# Donor-associated variation outweighs PMOS status in patient-derived iPSC transcriptomes: a genotyped, BMI-adjusted study of 14 lines

**DOI:** 10.64898/2026.11.09.765405

**Authors:** Masuma Khatun, Brayan Stiveen Rodriguez Morales, Vijayachitra Modhukur, Apostol Apostolov, Karolina Lundin, Riikka K. Arffman, Timo Tuuri, Juha S. Tapanainen, Terhi T. Piltonen, Andres Salumets

## Abstract

**Background:** Induced pluripotent stem cells (iPSCs) from patients have been little used to model polyendocrine metabolic ovarian syndrome (PMOS, formerly polycystic ovary syndrome, PCOS), and differences between unrelated donors can masquerade as disease effects. Earlier comparisons of three or four donors per group reported hundreds to thousands of differentially expressed genes. We asked whether a disease signature is detectable in a larger, genotyped panel, balanced for BMI category and BMI-adjusted, of full-phenotype patients.

**Methods:** We studied 14 clonal, transgene-free iPSC lines reprogrammed by CRISPR activation from endometrial stromal fibroblasts of seven women with PMOS meeting all three Rotterdam criteria, and seven controls frequency-matched for BMI category (<25 or >30 kg/m²). Bulk RNA-seq (16,489 expressed genes) was related to diagnosis, hormones, metabolic traits, age, and X-inactive specific transcript (XIST) status using gene-wise models, exact enumeration of all 3,432 group splits, and permutation-calibrated gene-set tests. Donors were genotyped within a cohort of 140 women and scored with an imputation-based genome-wide PMOS polygenic risk score (PRS). PMOS risk-locus genes were examined against a 202-line iPSC proteome.

**Results:** PMOS donors had higher anti-Müllerian hormone (AMH), testosterone and free androgen index than controls. No gene was differentially expressed between PMOS and control lines (false discovery rate < 0.1), and the number of nominally associated genes was no greater than for random 7-versus-7 splits of the lines (p = 0.75). This was unchanged with an alternative read quantification and with count-based models. With seven donors per group, the design could detect 1.8-fold differences per gene, compared with roughly 3-to 6-fold for the three-or four-donor designs used previously. Neither diagnosis nor any hormonal or metabolic trait explained more expression variance than chance, and neither PMOS GWAS-locus genes nor Hallmark pathways were enriched. Two control lines showed near-complete loss of XIST, potentially indicative of X-chromosome inactivation (XCI) erosion, and a third was borderline. Endocrine PMOS-locus genes (e.g., FSHR, AMH) were not expressed, whereas DNA-repair and growth genes were. Across 129 genotyped women, the PRS was higher in PMOS (odds ratio 1.81 per SD; AUC 0.65), but PMOS and control iPSC donors overlapped fully in polygenic risk.

**Conclusions:** In this panel of 14 undifferentiated iPSC lines from women with the full PMOS phenotype, no robust PMOS-associated transcriptomic differences were detected. Expression variation was dominated by line-to-line differences, consistent with donor background as their main source, although with one clone per donor a contribution of clonal variation cannot be excluded. Reduced XIST expression in some control lines is potentially indicative of, but does not establish, X-chromosome inactivation erosion. Disease-associated differences in differentiated, hormone-responsive cells cannot be excluded, and differences reported in smaller panels may partly reflect inter-donor variability. Future studies should use adequately sized panels, control for XIST erosion, select donors by polygenic risk, and examine hormone-responsive derivatives.

## Background

Polyendocrine metabolic ovarian syndrome (PMOS), formerly termed polycystic ovary syndrome (PCOS), is a common and heterogeneous endocrine-metabolic condition affecting reproductive, metabolic and psychological health across the lifespan. The nomenclature change followed a global consensus process intended to better reflect its broader endocrine and metabolic features (Stener-Victorin *et al*., 2026; Teede *et al*., 2026). Clinically, PMOS is characterised by combinations of hyperandrogenism, ovulatory dysfunction and polycystic ovarian morphology, as defined by the Rotterdam criteria (Fauser *et al*., 2004). Its aetiology is polygenic, involving reproductive hormone signalling, steroid regulation and metabolic pathways, with recent genome-wide studies expanding the number of associated loci (Day *et al*., 2018; Moolhuijsen *et al*., 2026). Nevertheless, how common risk variants influence specific human cell types and contribute to the heterogeneous clinical manifestations remains incompletely understood.

A major challenge in studying PMOS is distinguishing inherited susceptibility from the molecular changes acquired through the endocrine and metabolic environment. Disease-relevant tissues are often exposed to androgen, insulin, and gonadotropin signalling, making it difficult to separate genetic predisposition from secondary responses to the systemic milieu. The endometrium illustrates this complexity: endometrial stromal fibroblasts (eSF) from women with PCOS have been reported to exhibit inflammatory and oncogenic features, impaired progesterone-mediated decidualisation, and altered androgen responses (Piltonen *et al*., 2013, 2015; Khatun *et al*., 2017, 2021; Lavogina *et al*., 2021). Whether these abnormalities reflect stable, cell-intrinsic mechanisms or depend on the differentiated, hormone-responsive state remains unclear. Patient-derived induced pluripotent stem cells (iPSCs) offer a complementary approach to investigating inherited susceptibility outside the original endocrine environment. They provide a renewable pluripotent cell population that can be subsequently differentiated into disease-relevant lineages. However, reprogramming substantially remodels somatic cell identity and aspects of its epigenetic landscape. Consequently, disease-associated phenotypes in the tissue of origin may be attenuated or reset or emerge only after differentiation into an appropriate cell type. Thus, the absence of a disease-associated signal in undifferentiated iPSCs does not exclude genetic effects operating in hormone-responsive or lineage-restricted cells (Khatun *et al*., 2023, 2026b).

Interpretation of iPSC disease models also requires attention to variation unrelated to diagnosis. Studies of large human iPSC collections have identified donor genetic background as a major contributor to inter-individual molecular variation, with regulatory effects often depending on cell type and state (Rouhani *et al*., 2014; Kilpinen *et al*., 2017; Bonder *et al*., 2021). Clone, culture, and passage further contribute to heterogeneity. Female iPSC lines require particular attention to X-chromosome inactivation (XCI), which can erode during culture. Reduced expression of the X-inactive specific transcript (XIST) is commonly associated with XCI erosion and may accompany reactivation of X-linked genes and broader transcriptomic changes (Mekhoubad *et al*., 2012; Vallot *et al*., 2015; Raposo *et al*., 2025). The extent and consequences of this process can vary between lines and may persist following differentiation, including after genome editing (Thapa *et al*., 2026). Uneven distributions of these features between cases and control groups can therefore confound transcriptomic comparisons.

Previous iPSC studies have explored PMOS biology, but differences in donor numbers, source tissues, reprogramming approaches, and cellular models complicate comparisons. Two early studies examined undifferentiated iPSCs from three cases and three controls, reporting extensive transcriptional and metabolic differences (Min *et al*., 2018, 2019). Subsequent studies investigated iPSC-derived mesenchymal progenitor cells generated from luteinised granulosa cells (four cases and four controls) and iPSC-derived endothelial cells from three women with phenotype A PCOS and three controls (Huang *et al*., 2024; Wu *et al*., 2026). These studies suggest that disease-associated phenotypes may emerge in specific cellular contexts. However, their limited numbers of independent donors restrict the ability to distinguish diagnosis-associated effects from inter-individual variation, highlighting the importance of donor-level replication in iPSC transcriptomic studies (Germain and Testa, 2017).

Because PMOS is polygenic, clinical diagnosis is an imperfect proxy for inherited susceptibility: women meeting the same diagnostic criteria may carry substantially different burdens of common risk alleles, while unaffected women may also have considerable genetic liability (Joo *et al*., 2020; Moolhuijsen *et al*., 2026). Accordingly, iPSC studies of polygenic disorders increasingly use polygenic risk scores (PRS) to characterise genetic liability beyond clinical classification, sometimes selecting donors from opposite ends of the PRS distribution(Page *et al*., 2022; Sawada *et al*., 2024). However, individual PRS values require interpretation against an appropriately genotyped and processed reference population, and their association with disease should be evaluated in the target population before application to small donor panels (Choi *et al*., 2020). Complementary molecular resources can further contextualise genetic risk in pluripotent cells. iPSC expression quantitative trait locus (eQTL) maps identify genes whose expression is under genetic regulation in this cellular state (Kilpinen *et al*., 2017; Bonder *et al*., 2021) while population-scale proteomic data provide evidence of which expressed genes are translated into detectable proteins, given that mRNA abundance does not necessarily predict protein abundance (Liu *et al*., 2016; Mirauta *et al*., 2020).

We previously generated and characterised 14 clonal, transgene-free iPSC lines from eSF of seven women with PMOS meeting all three Rotterdam criteria and seven controls frequency-matched for body mass index (BMI) category, using CRISPR activation (CRISPRa)-based reprogramming (Khatun *et al*., 2026a). Here, we used this panel to investigate whether PMOS diagnosis is associated with a detectable transcriptomic signature in undifferentiated iPSCs and to assess the contribution of donor characteristics and cell-line features to gene-expression variability. We integrated RNA sequencing with clinical phenotypes and genome-wide genotype data to examine associations with diagnosis, hormonal and metabolic traits, age, and polygenic risk. The 14 iPSC donors were genotyped alongside 126 additional women from the same clinical population, yielding 129 individuals after quality control from the initial cohort of 140. This population-based resource enabled us to evaluate whether a genome-wide PMOS PRS derived from the largest GWAS to date (Moolhuijsen *et al*., 2026) captured genetic liability in this Finnish clinical population and whether clinical case–control selection had also generated a contrast in inherited risk between the iPSC groups. We additionally examined XIST expression as a potential indicator of XCI erosion and integrated expression data with PMOS-associated loci and the HipSci reference proteome of 202 iPSC lines (Mirauta *et al*., 2020) to assess which locus-associated genes were expressed and supported by detectable protein evidence in the pluripotent state. By defining the molecular features detectable in undifferentiated iPSCs, as well as the limitations of this cellular state, this study aims to inform the design and interpretation of iPSC-based models of complex endocrine disorders.

## Methods

### Participants and clinical phenotyping

Women were recruited at Oulu University Hospital. PMOS was diagnosed by the Rotterdam criteria (Rotterdam ESHRE/ASRM-Sponsored PCOS Consensus Workshop Group, 2004); all seven iPSC donors with PMOS were selected to meet all three criteria (oligo-/anovulation, clinical or biochemical hyperandrogenism, and polycystic ovarian morphology; Rotterdam phenotype A), the most complete and severe phenotype, so that the patient group was phenotypically homogeneous. Controls were frequency-matched to cases by BMI category rather than individually matched: each group comprised lean (BMI <25 kg/m²; 4 control, 3 PMOS) and obese (BMI >30 kg/m²; 3 control, 4 PMOS) donors, and no donor had a BMI of 25–30 kg/m². Because median BMI still differed between groups (31.1 vs 23.4 kg/m²; Table 1), BMI was additionally handled analytically by adjustment for BMI category and for continuous BMI, as recommended for frequency-matched designs (Pearce, 2016). Controls had regular cycles and no hyperandrogenism or PCOM. Fasting blood was analysed for AMH, total testosterone, DHT, SHBG, LH, glucose, insulin and HbA1c. FAI was calculated as 100 × testosterone (nmol/L)/SHBG (nmol/L), recalculated for all donors from the measured values, and HOMA-IR as glucose × insulin/22.5. Hirsutism was scored with the modified Ferriman–Gallwey (FG) score.

**Table 1.**
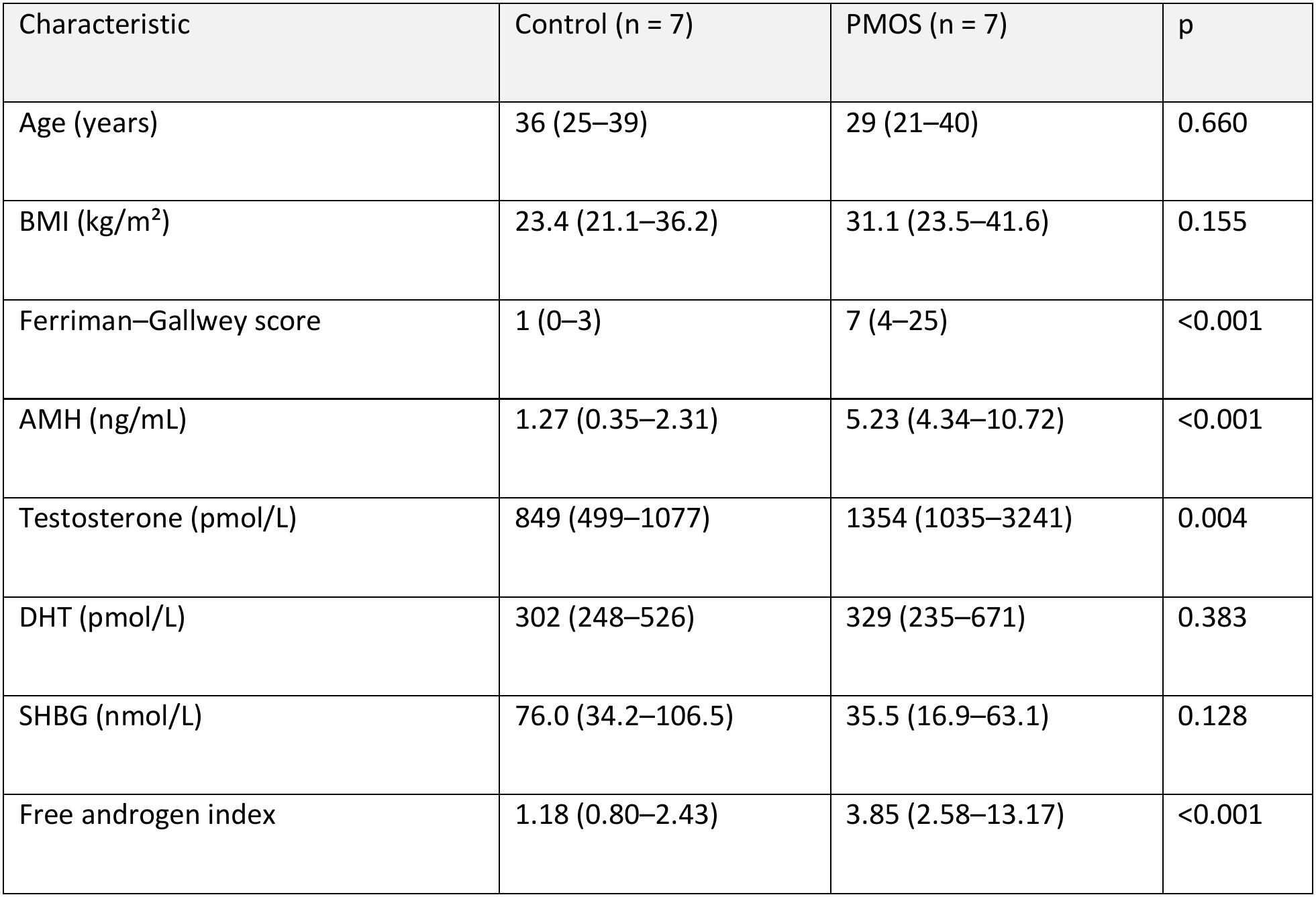
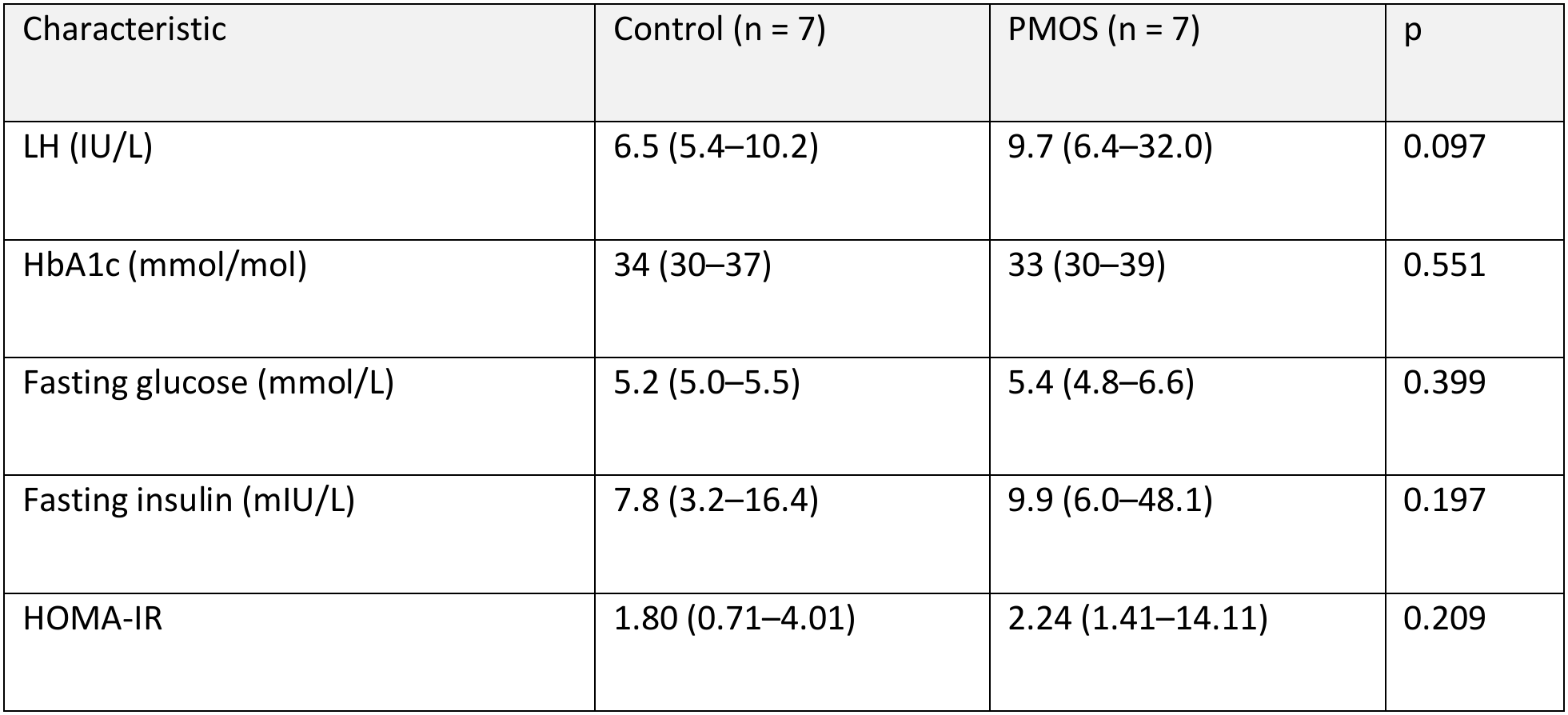
Clinical characteristics of the 14 iPSC donors. Median (range); two-sided Mann–Whitney U test (exact p-values, ties included). XIST-eroded lines (reverse-stranded counts): control 2/7, PMOS 0/7 (Fisher p = 0.46); including the borderline line HEL363.2, control 3/7 (p = 0.19; see Results).

| Characteristic | Control (n = 7) | PMOS (n = 7) | p |
|---|---|---|---|
| Age (years) | 36 (25–39) | 29 (21–40) | 0.660 |
| BMI (kg/m²) | 23.4 (21.1–36.2) | 31.1 (23.5–41.6) | 0.155 |
| Ferriman–Gallwey score | 1 (0–3) | 7 (4–25) | <0.001 |
| AMH (ng/mL) | 1.27 (0.35–2.31) | 5.23 (4.34–10.72) | <0.001 |
| Testosterone (pmol/L) | 849 (499–1077) | 1354 (1035–3241) | 0.004 |
| DHT (pmol/L) | 302 (248–526) | 329 (235–671) | 0.383 |
| SHBG (nmol/L) | 76.0 (34.2–106.5) | 35.5 (16.9–63.1) | 0.128 |
| Free androgen index | 1.18 (0.80–2.43) | 3.85 (2.58–13.17) | <0.001 |
| Characteristic | Control (n = 7) | PMOS (n = 7) | p |
| LH (IU/L) | 6.5 (5.4–10.2) | 9.7 (6.4–32.0) | 0.097 |
| HbA1c (mmol/mol) | 34 (30–37) | 33 (30–39) | 0.551 |
| Fasting glucose (mmol/L) | 5.2 (5.0–5.5) | 5.4 (4.8–6.6) | 0.399 |
| Fasting insulin (mIU/L) | 7.8 (3.2–16.4) | 9.9 (6.0–48.1) | 0.197 |
| HOMA-IR | 1.80 (0.71–4.01) | 2.24 (1.41–14.11) | 0.209 |

### iPSC derivation and culture

Generation and full characterisation of the 14 iPSC lines (hPSCreg UHi023-A to UHi036-A) have been described in detail (Khatun *et al*., 2026a). Briefly, eSF at passage 2 was reprogrammed with a CRISPRa-based method. Single colonies were picked at day 24, expanded clonally, and confirmed episome-free by passage 12. All lines expressed OCT4, NANOG, and TRA-1-60, differentiated into all three germ layers, and matched their donor eSF at 16 STR loci. Karyotypes were 46, XX except HEL350.3, which was mosaic for an isodicentric chromosome 20. Lines were maintained on Geltrex in Essential 8 medium and passaged with 0.5 mM EDTA (Khatun *et al*., 2026a).

### RNA sequencing

Total RNA was extracted from undifferentiated iPSCs at passages 15–17. Libraries were prepared with the TruSeq RNA Exome library preparation kit (Illumina) and sequenced single-end (80 cycles; the 14 libraries on one lane of one flow cell) on a NextSeq 1000 (Illumina), as described previously (Apostolov *et al*., 2025), yielding 15.3–27.8 million reads per line; the data confirmed reverse-strandedness (65.7% of reads assigned with reverse-strand and 5.3% with forward-strand counting; Additional file 1: Table S17). For the primary analysis, reads were quality-checked with FastQC, trimmed with Trim Galore and aligned to GRCh38 with HISAT2 in Galaxy (Apostolov et al., 2025), and gene-level counts were generated; these counts correspond to unstranded counting of the stranded library. To ensure that the results did not depend on this quantification, all analyses were repeated on an independent, fully specified quantification (STAR 2.7.10b, GRCh38, GENCODE v50, featureCounts 2.1.1; see Alternative quantification and count-based models), with the same transcriptome-wide conclusions. Gene-level counts were normalised by size factors (DESeq2 median-of-ratios; Love et al., 2014). Genes with ≥ 10 normalised counts in at least 7 of the 14 lines (n = 16,489) were analysed on the log2 (normalised count + 1) scale. XIST-eroded lines were defined as those with log2 XIST (normalised XIST count + 1) < 8 on the reverse-stranded counts (see Alternative quantification and count-based models). Because XIST overlaps the antisense TSIX transcript, unstranded counting of a stranded library splits reads between the two genes and underestimates XIST; HEL363.2, which was below the cut-off on the primary counts (7.25) but above it on the reverse-stranded counts (8.27), was treated as borderline and analysed as a sensitivity group.

### Alternative quantification and count-based models

To assess whether the results depended on read quantification or on the statistical model, the raw reads were also processed separately. Reads were aligned with STAR 2.7.10b (Dobin *et al*., 2013) to the GRCh38 primary assembly with GENCODE v50 annotation (Mudge *et al*., 2025) and counted with featureCounts 2.1.1 (Liao *et al*., 2014) as single-end, reverse-stranded reads (-s 2), with exon-level counts summarised to genes and multi-mapping reads excluded; unstranded counting (-s 0) was run as a check. Counts were normalised with DESeq2 median-of-ratios size factors (Love *et al*., 2014), and all transcriptome analyses were repeated on these counts with the same expression filter, models, permutation designs and random seeds. Sample identity was confirmed by correlating each line with every line of the primary count matrix. In addition, PMOS-versus-control differential expression was tested on raw counts with DESeq2 1.42 (negative binomial Wald test) and with limma-voom (limma 3.58) (Law *et al*., 2014; Ritchie *et al*., 2015) in R 4.3.3, with and without log2 XIST as a covariate. For HEL350.3, a chromosome 20 dosage profile was derived from per-gene z-scores across the 14 lines (log2 normalised counts of the reverse-stranded matrix, expressed genes), summarised as the median per chromosome arm (20p, genes ending before 25.6 Mb: 137 genes; 20q, genes starting after 30.4 Mb: 301 genes; GENCODE v50 coordinates) and compared with the other lines. An empirical p-value was obtained for each arm by comparing the HEL350.3 median with the medians of 1,000 random sets of the same number of expressed genes, drawn without replacement from the other chromosomes (seed 20; not matched for expression level); p = (number of random sets with an absolute median at least as large as the observed one + 1)/(1,000 + 1), so that the smallest attainable p is 0.001 (Additional file 1: Table S17).

### Genotyping and quality control

DNA from donor blood of 140 cohort women, including all 14 iPSC donors, was genotyped on the Illumina Global Screening Array v3 with the Estonian custom content (ESTchip-2_GSAv3-MD; 777,016 markers) at the Estonian Genome Centre core facility. Calls were made in GenomeStudio and exported with GRCh38 coordinates. Sample and variant QC used PLINK 1.9 and 2 (Chang *et al*., 2015) with standard thresholds (Anderson *et al*., 2010): sample call rate ≥ 0.98 (computed over autosomal variants with a variant call rate ≥ 95%); autosomal heterozygosity F within the mean ± 3 SD; genetic sex from X-chromosome F (female < 0.2); and KING-robust kinship (Manichaikul *et al*., 2010), removing one member of each first-degree pair (kinship > 0.177). Variants were kept if autosomal, with call rate ≥ 98%, MAF ≥ 0.01, and Hardy–Weinberg p ≥ 1 × 10⁻⁶. DNA contamination was assessed from the raw intensity data (B-allele frequency, BAF): the contaminating fraction was estimated as the slope of homozygous-call BAF against the frequency of the other allele, and samples with an estimate ≥ 9% were considered contaminated (five samples, 9.9-12.3%), and estimates of 3.5%-9% were flagged as possible low-level contamination (11 samples, 3.7-7.4%) and retained. Failure rates by BeadChip were compared by Fisher’s exact test. After QC, 129 of 140 samples and 447,260 variants were retained; a call-rate threshold of 0.97 was used as a sensitivity analysis. Line identity was checked independently by calling variants from the RNA-seq of each iPSC line at exonic array sites (STAR 2.7.10b, GRCh38, GENCODE v50; bcftools mpileup/call) and comparing them with all 140 donor genotypes. Genetic principal components were computed in PLINK 2 on the array genotypes of the 129 QC-passing women (autosomal SNPs with MAF ≥ 0.05, long-range LD regions such as the MHC and the 8p23 and 17q21 inversions excluded, LD pruning with a 200-SNP window and r² 0.2; 78,749 SNPs). Lead variants of Moolhuijsen et al. (2026) were matched to the array by rsID and, after imputation, to the imputed data. Alleles were aligned to the reported risk allele on both strands, and risk-allele frequencies were compared with the GWAS.

### Reference iPSC proteome and QTL data

Protein was not measured in the 14 lines. Instead, we used the publicly available population-scale reference proteome of the Human Induced Pluripotent Stem Cell Initiative (HipSci), in which protein abundance was quantified by tandem mass tag (TMT) mass spectrometry in 202 iPSC lines from 151 unrelated donors with matched RNA-seq and genotypes, and 654 protein QTLs (pQTLs) were mapped (Mirauta *et al*., 2020). The data were used for three purposes: (i) to determine whether genes expressed in our lines, and in particular the PMOS-locus genes, are translated into detectable protein in undifferentiated iPSCs; (ii) to estimate, for each gene, how closely mRNA tracks protein across iPSC lines (reference RNA–protein correlation), and hence how likely an mRNA difference in our lines is to be reflected at the protein level (Liu *et al*., 2016); and (iii) to identify PMOS-locus genes under genetic control (cis-eQTL or pQTL) in iPSCs. All values were taken from the supplementary source data of (Mirauta *et al*., 2020); gene matching, filtering, and enrichment analyses are described below (Protein-level cross-reference).

## Statistical analysis

### Group comparison of donors

two-sided Mann–Whitney U tests with exact p-values, obtained by enumerating all possible assignments of the donors to the groups (3,432 for 7 versus 7; tied values included).

#### Gene-wise models

for each gene, log2 expression was regressed on PMOS status by ordinary least squares, with and without log2 XIST as a covariate. P values were adjusted by Benjamini–Hochberg (FDR).

#### Transcriptome-wide signal

for each trait, we counted the genes with nominal p < 0.05 (Spearman correlation). For PMOS status, this count was compared with the exact null from all 3,432 possible 7-versus-7 splits of the 14 lines. For continuous traits, it was compared with 2,000 permutations of trait values. This tests whether a trait explains more expression variation than chance while respecting gene– gene correlation. For PMOS status, the test was repeated on subsets of lines (without HEL350.3 and HEL351.3, without HEL350.3 only, and without the two XIST-eroded lines, with and without the borderline line HEL363.2), each against its own exact null (Additional file 1: Table S7).

#### Principal components

computed on the 1,000 most variable genes and correlated (Spearman) with donor traits; PCA was repeated on all 16,489 expressed genes.

#### Sample correlation

Pearson correlations between lines were computed on all 16,489 expressed genes (log2 scale) and clustered by average linkage. Whether lines correlated more strongly within than between groups (diagnosis; BMI <25 vs <30; XIST-eroded vs XIST-expressing) was tested by the difference between mean within-group and between-group r, compared with all possible splits of the 14 lines.

#### Variance partition

each gene’s log2 expression was modelled on PMOS status, BMI, age and log2 XIST together. The variance uniquely explained by each factor was its Type II sum of squares as a fraction of the total; the remainder is residual. With 14 lines, every factor explains a few percent of variance by chance, so the four covariates were permuted jointly across lines (1,000 permutations), and each factor’s median was compared with its null distribution. The model without XIST, as used in an earlier analysis of these data, is reported for comparison.

#### BMI adjustment

the gene-wise PMOS model was refitted with BMI category (see Participants) and, as sensitivity, with continuous BMI. A PMOS × BMI-category interaction model was fitted for completeness. Donor characteristics were also compared within BMI strata by two-sided Mann–Whitney U tests with exact p-values, obtained by enumerating all 35 possible assignments of3 versus 4 donors (tied values included); without ties, the smallest attainable two-sided p is 0.057 (Additional file 1: Table S11).

#### Protein-level cross-reference

protein abundances (proteins recurrently detected in at least 30 lines; Figure 1—source data 3 of Mirauta et al., 2020), RNA expression (source data 2) and genes with a cis pQTL or eQTL at FDR < 10% (Figure 2—source data 1–2) were taken from the original supplementary data. Our gene symbols were mapped to Ensembl gene identifiers through the HGNC protein-coding gene set (current and unique previous symbols; accessed 4 Nov 2026) (Seal *et al*., 2026), and HipSci proteins were matched by Ensembl identifier; where a gene mapped to several protein groups, the group with the most peptides was used. For each gene, the reference RNA–protein correlation was the Pearson correlation between HipSci RNA and log10 protein abundance across the 201 lines with both measurements. We recorded protein detection for all 16,489 expressed genes, reported for all genes and for the 13,241 protein-coding genes, and for the PMOS-locus genes, and listed as protein-supported candidates the genes with nominal PMOS-versus-control p < 0.05, detected protein, and a reference RNA–protein correlation r ≥ 0.3. Functional enrichment (over-representation) analysis was performed with g:Profiler (Kolberg *et al*., 2023) (GO biological process, molecular function and cellular component, KEGG and Reactome; g:SCS-corrected p < 0.05), each gene set against a matched custom background: genes with and without detected protein against all expressed protein-coding genes, and the protein-supported candidates against all genes passing the same protein and correlation filters. All code is available in the GitHub repository and the Zenodo archive (see Additional file 3 for the summary of methods and the map from figures and tables to scripts).

**Fig. 1.**
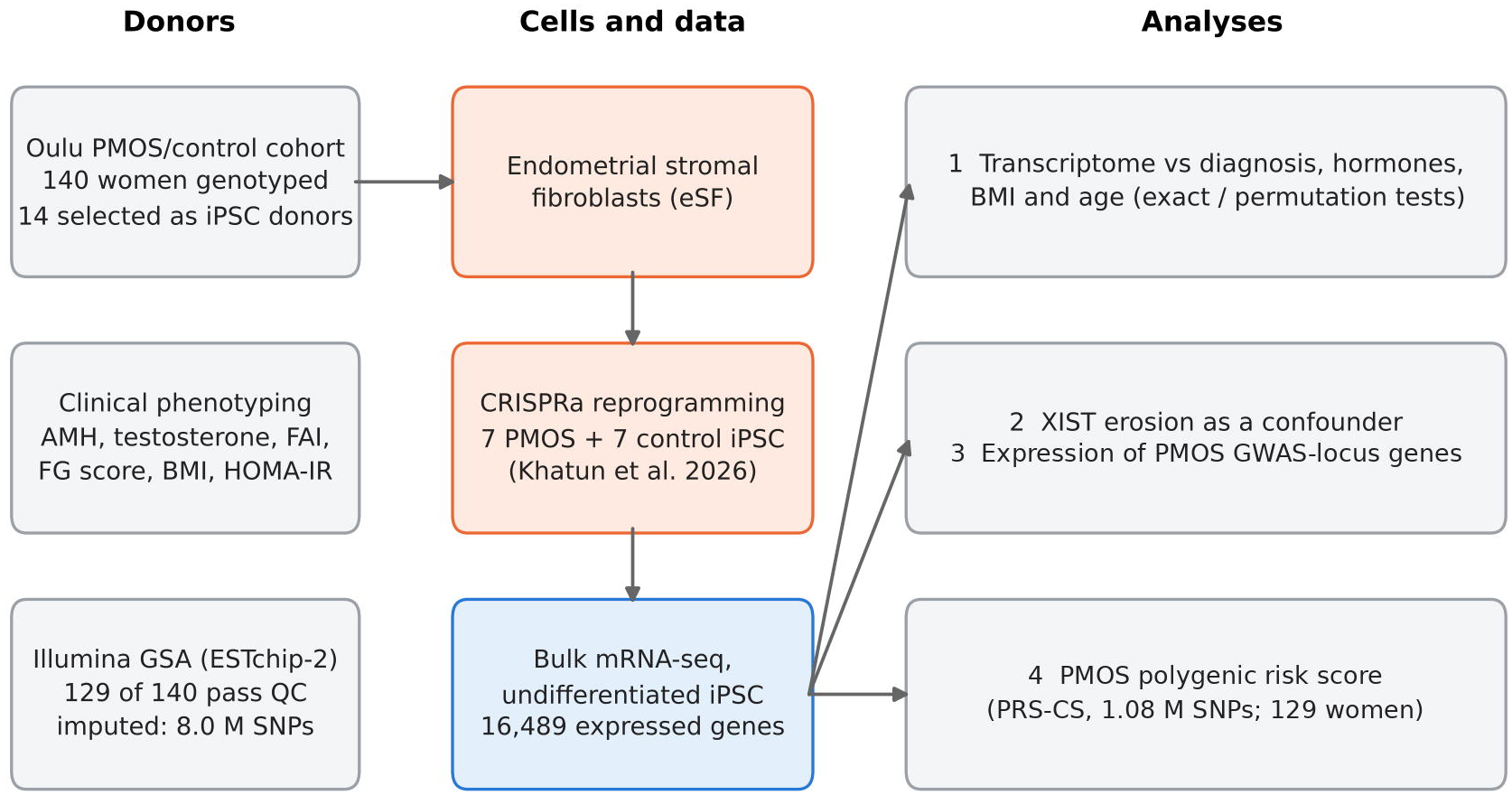
Study design. Fourteen clonal iPSC lines were derived by CRISPRa reprogramming of endometrial stromal fibroblasts from 7 women with PMOS, all meeting all three Rotterdam criteria (phenotype A), and 7 controls frequency-matched for BMI category (<25 or >30 kg/m²) (Khatun et al., 2026a). The donors were drawn from an Oulu cohort of 140 women genotyped on the Illumina GSA. Bulk mRNA-seq of undifferentiated lines was integrated with donor phenotypes and genotypes. Genotypes were imputed to the TOPMed r3 reference, and a genome-wide PMOS polygenic risk score was calculated for the 129 women passing genotype QC.

**Fig. 2.**
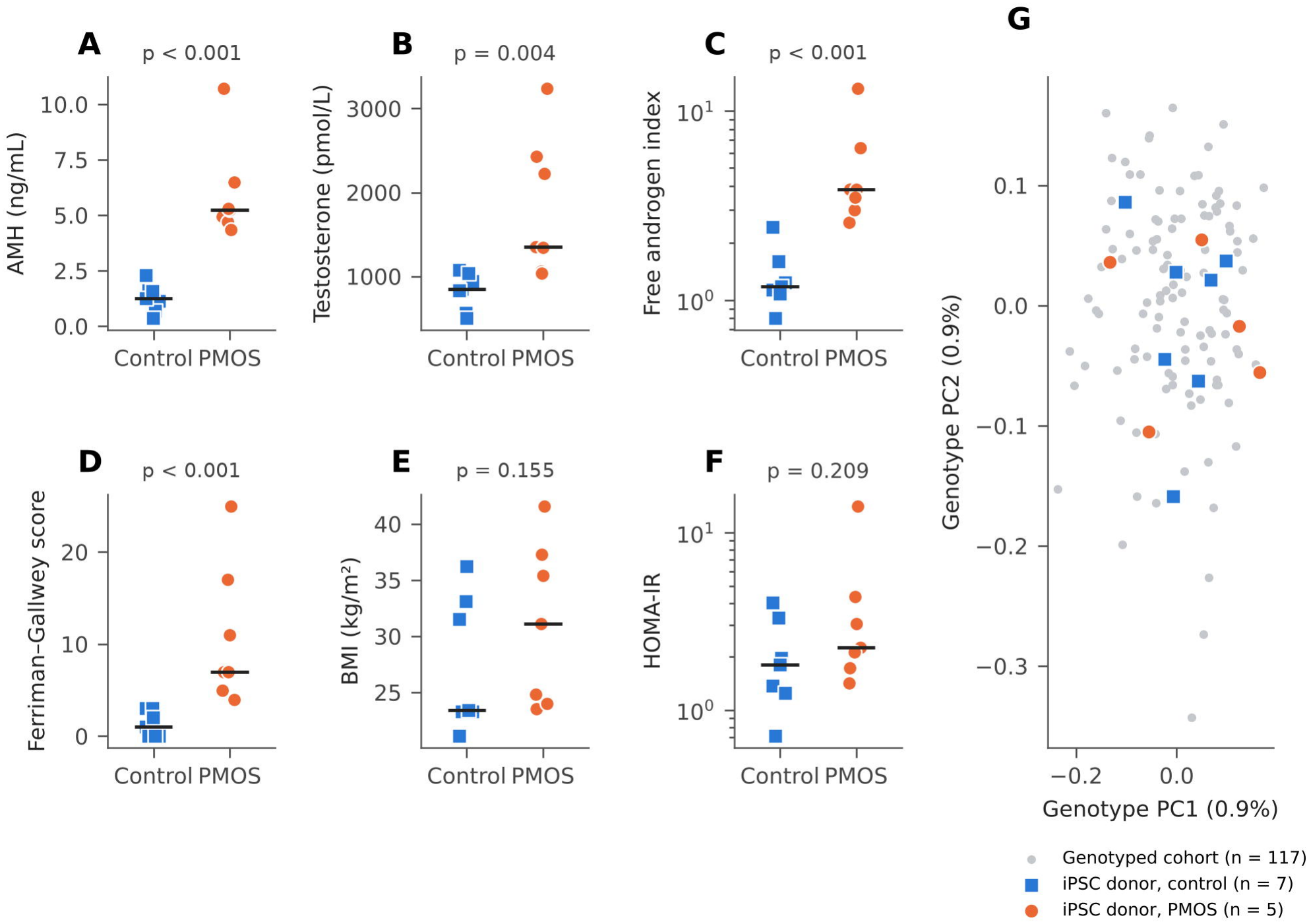
Clinical and genetic characteristics of the iPSC donors. (A–F) AMH, testosterone, free androgen index (log scale), Ferriman–Gallwey score, BMI and HOMA-IR (log scale) in control (blue squares) and PMOS (orange circles) donors. Bars, medians; p, two-sided Mann–Whitney U test (exact, ties included). (G) Genotype principal components 1 and 2 of the 129 QC-passing women (grey), with the 12 QC-passing iPSC donors highlighted; HEL351.3 and HEL360.2 failed genotype QC and are not shown. Each component explains < 1% of the variance, consistent with a genetically homogeneous cohort.

#### Literature candidate genes

60 genes previously linked to PMOS by GWAS or pathobiology, compiled for an earlier analysis of this dataset, were looked up in the gene-wise results.

#### PMOS GWAS genes

consensus genes for the 29 loci were taken from Moolhuijsen *et al*., 2026 (33 genes; four loci contribute two genes each); the “CYP3 complex” locus (rs148982377), for which the paper names no gene, was assigned to CYP3A7, named in its description, and CYP3A5. A gene-set score (mean of gene-wise z-scores of expressed genes) was compared with diagnosis and traits. Significance came from 10,000 random gene sets matched on expression level (20 bins).

*Gene-set enrichment:* preranked gene-set enrichment analysis (fgsea 1.28, fgseaMultilevel; minimum and maximum set sizes 5 and 500; (Korotkevich *et al*., 2021)) was run for the GWAS consensus genes, the 60 literature genes, and the MSigDB Hallmark sets ((Liberzon *et al*., 2015); release 2025.1, msigdbr 25.1.1) on all 16,495 genes expressed in the alternative counts, ranked by the DESeq2 Wald statistic for PMOS versus control; the OLS model, the XIST-adjusted model and the model without HEL350.3 were used as alternative rankings. Because gene-permutation p-values assume independent genes and are too optimistic when co-expressed genes are tested in 14 samples, significance was calibrated by exact sample-label permutation over all 3,432 group splits (1,716 without HEL350.3).

#### Published PMOS iPSC signature

the full gene lists of Min et al. (2018, 2019) are not public; their 20 named top genes (10 per direction) were therefore looked up in the gene-wise results, and the proportion changing in the same direction was compared with chance (binomial test, descriptive).

#### Imputation and PMOS polygenic risk score

Genotypes were phased and imputed on the TOPMed Imputation Server (reference panel r3, GRCh38; Eagle phasing, Minimac4) (Das *et al*., 2016; Taliun *et al*., 2021). Imputed SNPs were kept if biallelic, with imputation R² ≥ 0.3, MAF ≥ 0.01, and Hardy–Weinberg p ≥ 1 × 10⁻⁶, and genotyped SNPs if their leave-one-out empirical R² was ≥ 0.5, leaving 8,032,962 SNPs (435,979 genotyped and 7,596,983 imputed; median imputation R² 0.983). Because the GWAS publication does not provide PRS weights and the full three-strata summary statistics are not public, posterior SNP weights were derived with PRS-CS (Ge *et al*., 2019) from the public European-ancestry, age-adjusted summary statistics excluding 23andMe (GWAS Catalog GCST90570579; n = 420,703) (Moolhuijsen *et al*., 2026), with φ = 10⁻⁴, the median per-SNP sample size (412,081), a = 1, b = 0.5 and 1,000 MCMC iterations. The main score used the UK Biobank European LD reference panel (HapMap3 SNPs), restricted to SNPs present in the imputed data (1,079,002 SNPs, 96.8% of the panel; 15.2% before imputation). Scores were calculated from imputed dosages in PLINK 2 (Chang *et al*., 2015) and standardised within the 129 women passing QC. Sensitivity analyses used the 1000 Genomes European LD panel, weights for all panel SNPs, only SNPs with imputation R² ≥ 0.8, and two alternative sample sets (133 women at call rate ≥ 0.97 without two further contaminated samples; 128 women without one borderline-contaminated sample). A 29-variant genetic risk score (GRS29) was calculated from the risk alleles and log odds ratios of the lead variants (Moolhuijsen et al., 2026); strand-ambiguous variants were oriented by allele frequency. PRS was compared between PMOS cases and controls by the Wilcoxon rank-sum test and by logistic regression on the standardised score (odds ratio per SD), with discrimination summarised as the area under the ROC curve (AUC) with DeLong 95% CI. These comparisons were not adjusted for genetic principal components, age, or BMI. Scores were verified by independent recomputation from the imputed dosages for one chromosome (PRS) and for the GRS29 (maximum difference < 2 × 10⁻⁵).

#### Detectable effect size

To compare panel sizes, each gene’s between-line standard deviation (SD) was derived from the gene-wise PMOS-versus-control model (SD = |log2 fold change| / |t| / √(2/7); 15,562 genes with p < 0.95). With the median SD (0.29 log2 units; interquartile range 0.19–0.47), the smallest log2 fold change detectable with 80% power by a two-sided two-sample t-test was calculated from the noncentral t distribution for 3, 4, 7, 10, 15, and 20 single-clone donors per group, at per-gene α = 0.05 and α = 0.001 (Additional file 1: Table S16).

Analyses used Python 3.13 (NumPy, SciPy, pandas, Matplotlib); genotype QC, imputation post-processing, and PRS used PLINK 1.9/2, bcftools, and PRS-CS on the University of Tartu high-performance computing cluster. The alternative quantification used STAR 2.7.10b and featureCounts 2.1.1, and the count-based models R 4.3.3 (DESeq2 1.42, limma 3.58). Gene-set enrichment used R 4.3 with fgsea 1.28 and msigdbr 25.1.1. The analysis code is available at GitHub (https://github.com/BrayanRodriguez2/PMOS-iPSC-study-analysis-code) and archived at Zenodo (https://doi.org/10.5281/zenodo.24025021).

## Results

### PMOS donors differ in reproductive hormones but not in metabolic traits

As expected for phenotype A, PMOS donors had higher AMH (median 5.23 vs 1.27 ng/mL, p < 0.001), testosterone (1,354 vs 849 pmol/L, p = 0.004), FAI (3.85 vs 1.18, p = <0.001), and FG score (7 vs 1, p = <0.001) than controls (Table 1, Fig. 2A–D). BMI (31.1 vs 23.4 kg/m², p = 0.16), HOMA-IR (2.24 vs 1.80, p = 0.21), HbA1c, glucose, and age did not differ significantly (Fig. 2E–F). BMI ranged widely in both groups (control 21.1–36.2, PMOS 23.5–41.6 kg/m²). One PMOS donor (HEL352.2) had marked insulin resistance (HOMA-IR 14.1).

Of the 140 genotyped women, 129 passed genotype QC (Additional file 1: Table S2). All samples were genetically female. Eleven samples were excluded: five showed the B-allele-frequency pattern of a DNA mixture, with an estimated 10–12% of a second genome (all five also had call rates below 0.98 and three had excess heterozygosity); one had reduced heterozygosity (excess homozygosity, F = 0.081); one was the first-degree relative of another participant (kinship 0.28); and four had borderline call rates just below the 0.98 threshold (0.9749–0.9785) without other abnormalities. QC failures clustered on one of seven BeadChips (5 of its 23 samples vs 6 of 117 on the other chips; Fisher p = 0.018), consistent with a handling event during genotyping rather than with donor characteristics. Two PMOS donors, HEL351.3 and HEL360.2, were among the contaminated samples and were excluded from the PRS analyses; eleven retained samples, including that of control donor HEL362.1, had a low-level contamination estimate (3.7– 7.4%); because this estimator is not yet validated, these samples were flagged but kept. Variants called from the RNA-seq of each line matched the recorded donor for all 14 lines (concordance 0.976–0.982, against at most 0.811 with any other donor), so the contamination affected the donor DNA on the array, not the identity of the lines. The cohort was genetically homogeneous: each of the first five genetic principal components explained < 1% of the variance (0.86–0.92%), so none reflected population structure, and all 12 QC-passing iPSC donors lay within the cohort distribution (Fig. 2G).

### The iPSC transcriptome does not separate PMOS from control lines

Of 16,489 expressed genes, none differed between PMOS and control lines at FDR < 0.1 (Fig. 3C; Additional file 1: Table S4), and adding XIST as a covariate did not change this. In the gene-wise regression, only 443 genes reached nominal p < 0.05, fewer than the 824 expected by chance. Exact enumeration of all 3,432 possible group splits confirmed that diagnosis explained no more variation than a random split (observed 505 genes by Spearman correlation, null median 708, p = 0.75; Fig. 3D). The most nominally associated genes (TMEM86B, LMLN, C11orf54, INSYN2A, ZNF286B; Fig. 3C; Additional file 2: Fig. S2; Additional file 1: Table S14) overlapped with the top candidates of an earlier analysis of this dataset, but none approached FDR significance. The ranking of these nominal genes was, however, not robust to quantification: with the alternative quantification, only 25 of the top 40 genes remained among the top 40, so Fig. S2 and Table S14 are illustrative and do not represent a disease signature.

**Fig. 3.**
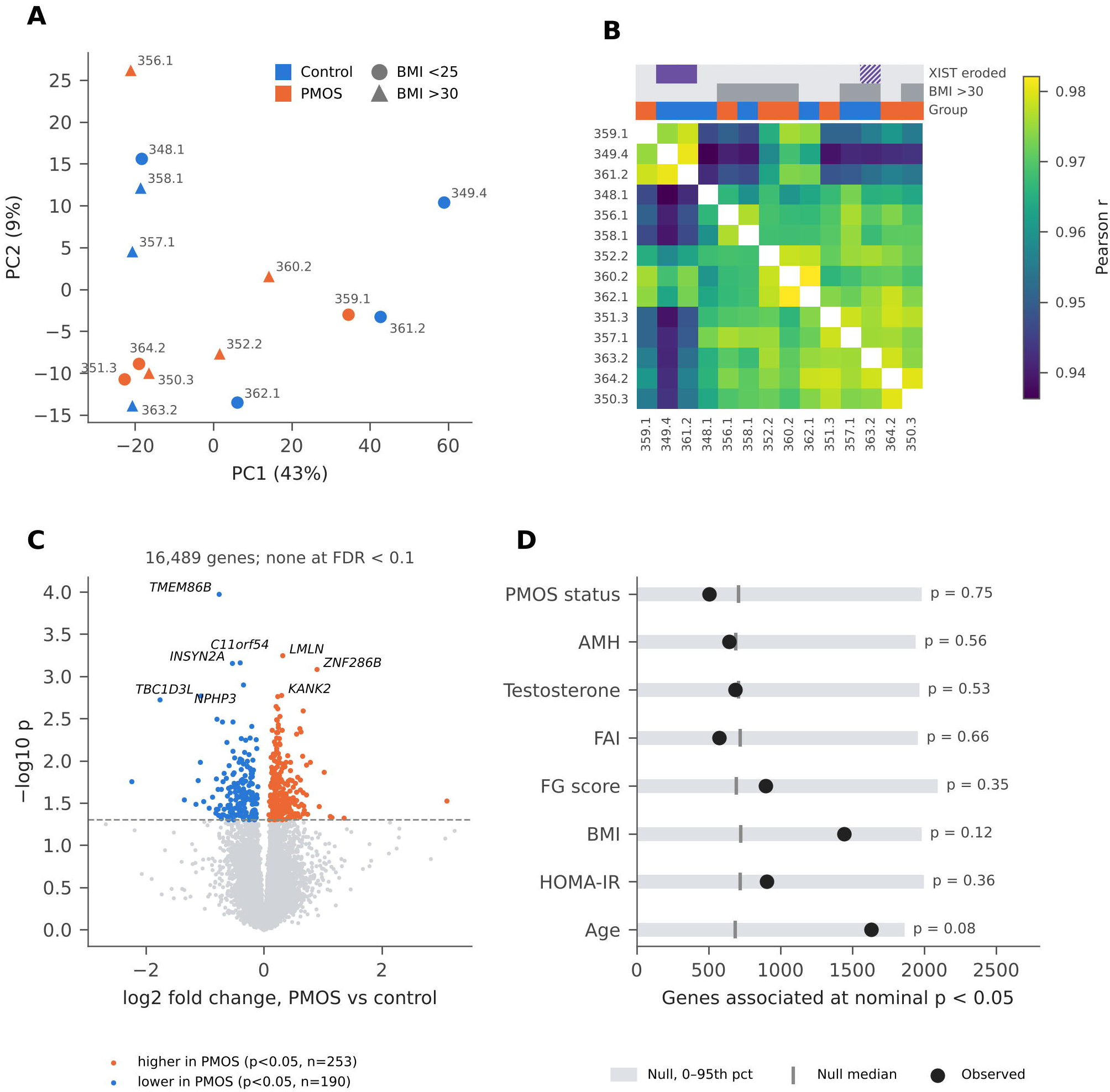
The iPSC transcriptome does not separate PMOS from control lines. (A) PCA of the 1,000 most variable genes; colour, diagnosis; shape, BMI category (circle <25, triangle >30 kg/m²); labels are line numbers. (B) Pearson correlation between lines (16,489 genes, log2), ordered by average-linkage clustering; bars show diagnosis, BMI >30 and XIST erosion (hatched, HEL363.2 borderline). Within-versus between-group correlation, exact tests: diagnosis p = 0.79, BMI p = 0.08, XIST erosion p = 0.011 (two eroded lines; 0.033 with the borderline HEL363.2 included). (C) Volcano plot of PMOS versus control (16,489 genes); coloured, nominal p < 0.05; dashed line, p = 0.05; no gene reaches FDR < 0.1. (D) Genes associated with each trait at nominal p < 0.05 (black dots) against the null distribution (grey bar, 0–95th percentile; tick, median). The null for PMOS status uses all 3,432 7-vs-7 splits; for traits, 2,000 permutations. p, permutation p-value.

Principal component analysis showed no group separation (Fig. 3A). PC1 (43% of variance) showed only a nominal correlation with donor age (ρ = −0.54, p = 0.044; p = 0.095 with the alternative quantification; not significant given the 40 trait–component tests; Additional file 2: Fig. S1B). No other trait was associated with PC1–PC5 at p < 0.05. PCA on all 16,489 genes gave the same picture (PC1 35% of variance; Additional file 2: Fig. S1A).

Sample-to-sample correlation was uniformly high (Pearson r = 0.936–0.982; Fig. 3B; Additional file 1: Table S8). Lines were not more correlated within than between diagnostic groups (mean within-group r 0.964 vs between-group 0.966; exact p = 0.79), and only weakly by BMI category (p = 0.08). The XIST-eroded lines were more similar to each other than expected (exact p = 0.011 over 91 splits; 0.011 with the alternative quantification), and the result was weaker when the borderline line HEL363.2 was also counted (p = 0.033 over 364 splits; 0.030 with the alternative quantification): HEL349.4 and HEL361.2 clustered with HEL359.1, the three lines with the highest PC1 scores.

The null result was robust. Excluding HEL350.3 (mosaic idic(20)) and HEL351.3 (contaminated array DNA) gave 491 nominal genes against a null median of 625 (exact p = 0.66, 792 splits). Excluding the two XIST-eroded lines gave 600 against 679 (exact p = 0.60, 792 splits), and also excluding the borderline line HEL363.2 gave 642 against 580 (p = 0.45, 330 splits). No gene reached FDR < 0.1 in either analysis (Additional file 1: Table S7).

The results were also robust to read quantification and to the statistical model (Additional file 1: Table S17). All 14 libraries were of good and homogeneous quality (15.3–27.8 million reads; 88.2–90.2% uniquely mapped; 77.1–80.2% assigned to genes; 6.2–9.0% mitochondrial reads; DESeq2 size factors 0.70–1.36), and each line correlated best with the corresponding line of the primary count matrix (r = 0.965–0.970; 14 of 14 lines), confirming the sample labels. With reverse-stranded counts, 422 genes reached nominal p < 0.05 (443 with the primary counts) and none reached FDR < 0.1 (634 and none with XIST as a covariate); log2 fold changes correlated closely with the primary estimates (r = 0.94). DESeq2 and limma-voom on raw counts gave the same answer (435 and 360 nominal genes; none at FDR < 0.1; rank correlation with the gene-wise regression 0.97 and 0.99), also with XIST as a covariate (739 and 529 nominal genes; none at FDR < 0.1). The exact-split test (497 genes vs null median 701; p = 0.74), line correlations (r = 0.932–0.983), principal components, variance partition, BMI-adjusted model (438 nominal genes, 342 shared) and sensitivity analyses (Table S7) gave consistent results. Differences between the two quantifications arose from read counting (strandedness and handling of multi-mapping reads).

The RNA-seq data also supported the karyotype of HEL350.3, which is mosaic for an isodicentric chromosome 20 (Khatun *et al*., 2026a). In this line, genes on 20q were expressed above (median z +1.44; highest of the 14 lines) and genes on 20p below (median z −1.87; lowest of the 14 lines) the other lines, with the switch at the centromere, whereas the arm medians of the other 13 lines lay between −0.48 and +0.33 (Additional file 1: Table S17). Without HEL350.3, 343 genes reached nominal p < 0.05, and none reached FDR < 0.1 (log2 fold-change correlation with the full model, r = 0.95), so this line does not drive the main result. Chromosome 20 genes of HEL350.3 should nevertheless be interpreted with caution.

The panel was also designed to detect smaller effects than earlier PMOS iPSC designs. Given the between-line variability observed here (median SD 0.29 log2 units per gene), seven donors per group detect a 1.8-fold difference with 80% power at a per-gene α of 0.001, whereas four and three donors per group would require 2.9-fold and 5.9-fold differences (at α = 0.05: 1.39-, 1.62- and 1.87-fold; Additional file 1: Table S16). A two-fold PMOS effect on a gene would therefore have been detectable in this panel but not in a three-or four-donor design.

### Donor hormones and metabolic traits leave no transcriptome-wide trace

We next asked whether continuous donor traits explained expression better than the binary diagnosis (Fig. 3D; Additional file 1: Table S5). AMH (p = 0.56), testosterone (p = 0.53), FAI (p = 0.66), FG score (p = 0.35) and HOMA-IR (p = 0.36) did not exceed their permutation nulls. BMI (1,442 genes vs null median 725, p = 0.12) and age (1,632 vs 685, p = 0.08) showed the largest, non-significant excess. No single gene was associated with any trait at FDR < 0.1.

### Diagnosis, BMI and XIST each explain no more variance than chance; age shows a borderline excess

Modelling each gene on PMOS status, BMI, age and XIST together, PMOS status uniquely explained a median of 3.2% of expression variance, BMI 5.2%, age 7.5% and XIST 6.2%, leaving 64% residual (Fig. 4A; Additional file 1: Table S9). With 14 lines, any single factor explains about 4% per gene by chance (permutation null medians 3.8–3.9%). Against this null, PMOS status (p = 0.72), BMI (p = 0.22) and XIST (p = 0.11) did not exceed chance, whereas age showed a borderline excess (p = 0.04; 0.053 with the alternative quantification), in line with its nominal correlation with PC1. Without XIST in the model, the medians were 3.0% (PMOS), 6.3% (BMI) and 7.7% (age), with 74% residual, close to an earlier analysis of these data; the larger shares for BMI and age reported are therefore mostly within chance expectation.

**Fig. 4.**
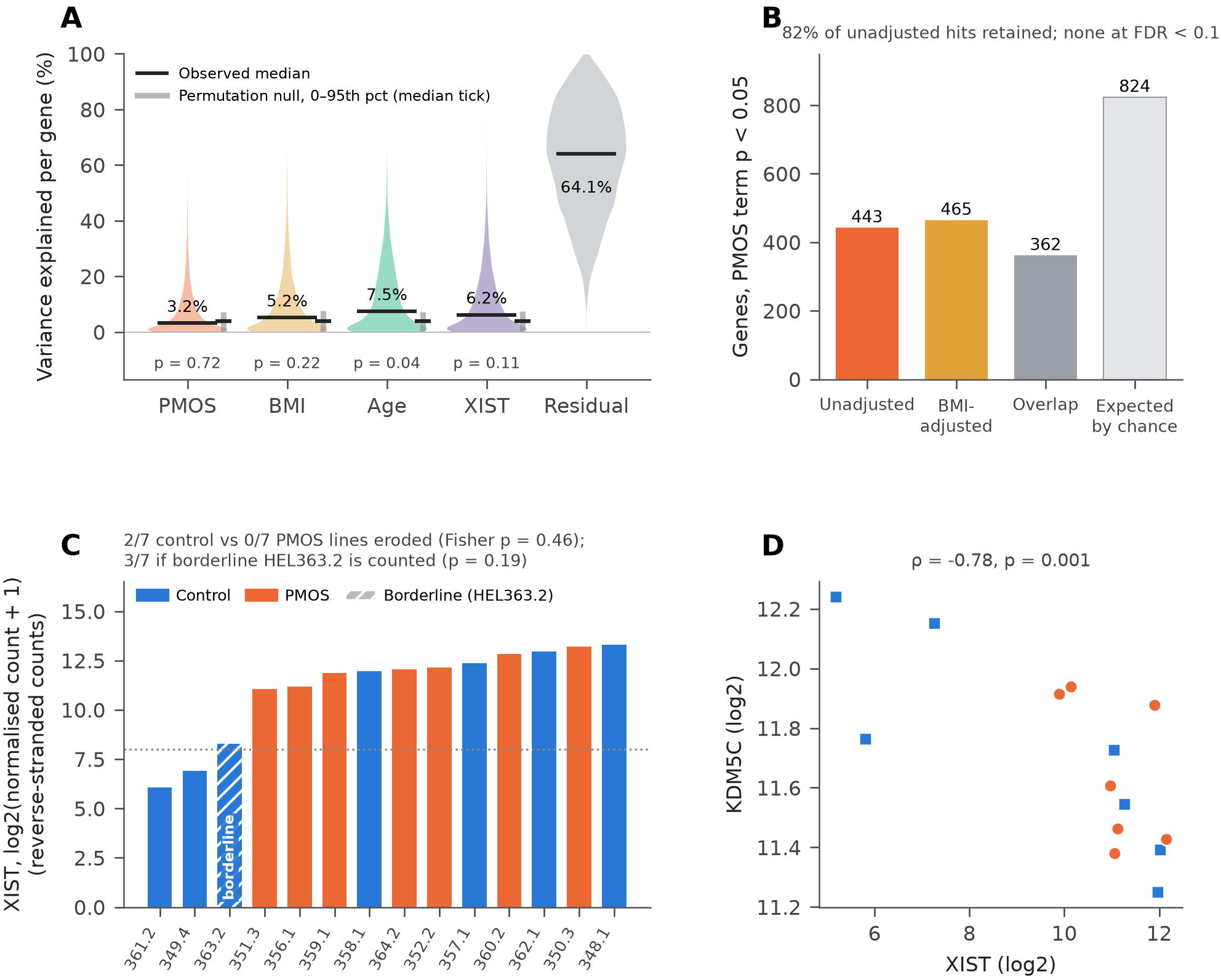
Sources of expression variation, BMI adjustment, and XIST erosion. (A) Variance partition: per-gene share of expression variance uniquely explained by PMOS status, BMI, age, and XIST (Type II sum of squares / total), and residual. Black line, observed median (value shown); grey bar and tick, 0–95th percentile and median of 1,000 joint permutations of the covariates; p, permutation p-value. (B) Genes with PMOS-term p < 0.05 without and with adjustment for BMI category, their overlap, and the number expected by chance. (C) XIST expression per line (reverse-stranded counts), ordered; the dotted line at log2 = 8 separates eroded from XIST-expressing lines. HEL349.4 and HEL361.2 lie below the line; HEL363.2 (labelled) is borderline (log2 8.27 with reverse-stranded counting; 7.25 with the primary counts). Both clearly eroded lines are controls (2/7 vs 0/7, Fisher p = 0.46; 3/7 vs 0/7, Fisher p = 0.19, if HEL363.2 is included). (D) XIST versus the X-linked gene KDM5C (primary counts); Spearman ρ.

Adjusting the PMOS comparison for BMI category changed the nominal gene list only at the margins: 465 genes reached p < 0.05 (443 without adjustment; 824 expected by chance), 362 (82%) of the unadjusted genes were retained, and none reached FDR < 0.1 (Fig. 4B; Additional file 1: Table S10). Continuous BMI gave similar results (487 genes, none at FDR < 0.1), and no gene showed a PMOS × BMI interaction at FDR < 0.1. Within both BMI strata, PMOS donors had higher AMH, testosterone, FAI and Ferriman–Gallwey score than controls, with complete separation of the groups (exact p = 0.057 each, the smallest p attainable with 3 versus 4 donors); SHBG was lower in PMOS donors in the lean stratum only (Additional file 1: Table S11).

### Reduced XIST expression, potentially indicativeof XCI erosion, is observed only in control lines

XIST expression was bimodal. Two lines, HEL361.2 and HEL349.4, showednear-complete loss of XIST with every quantification (35 and 55 normalised counts, log2 5.2 and 5.8 on the primary counts; log2 6.1 and 6.9 on reverse-stranded counts), potentially indicative of XCI erosion, whereas the other 11 lines, excluding the borderline line, expressed 949–4,520 counts (Fig. 4C). HEL363.2 was borderline: with reverse-stranded counting, which does not split reads with the overlapping antisense TSIX transcript, its XIST level (log2 8.27) lay just above the cut-off and clearly above those of the other two lines (7.25 on the primary counts, below the cut-off), suggesting partial erosion, for example, a mixture of eroded and non-eroded cells in the bulk culture. Both clearly eroded lines were controls (2/7 vs 0/7; Fisher p = 0.46; 3/7 vs 0/7, p = 0.19, if HEL363.2 is counted; Fig. 4C). XIST loss was accompanied by higher expression of the X-linked demethylase KDM5C (ρ = −0.78, p = 0.001; Fig. 4D); ρ = −0.81 with the alternative quantification and of CTPS2 (ρ = −0.89, p < 0.001), suggestive of X-chromosome reactivation. Adjusting for XIST shifted the PMOS-versus-control p-value distribution towards smaller values (627 vs 443 genes at p < 0.05; Additional file 2: Fig. S1C), suggesting that XIST-associated variation may partly mask group differences. Even so, no gene reached FDR < 0.1.

### Endocrine PMOS-locus genes are silent in iPSCs, DNA-repair and growth genes are highly expressed

Of the 33 consensus genes at the 29 PMOS loci (Moolhuijsen et al., 2026), 24 were expressed (Fig. 5A; Additional file 1: Table S6). The genes most closely linked to the endocrine phenotype were not expressed: FSHR, FSHB, AMH, SHBG, INHBB, GATA4, TOPAZ1, CYP3A7 and HHAT. In contrast, DNA-repair and genome-maintenance genes were highly expressed, including MSH6 (98th percentile of expressed genes) and CHEK2 (85th), as were NEIL2, FANCC, HELB and RAD50. So were growth and signalling genes: HMGA2 (97th), MYO10 (96th), MAPRE1 (95th), YAP1 (94th), ERBB3 (91st), THADA and DENND1A.

**Fig. 5.**
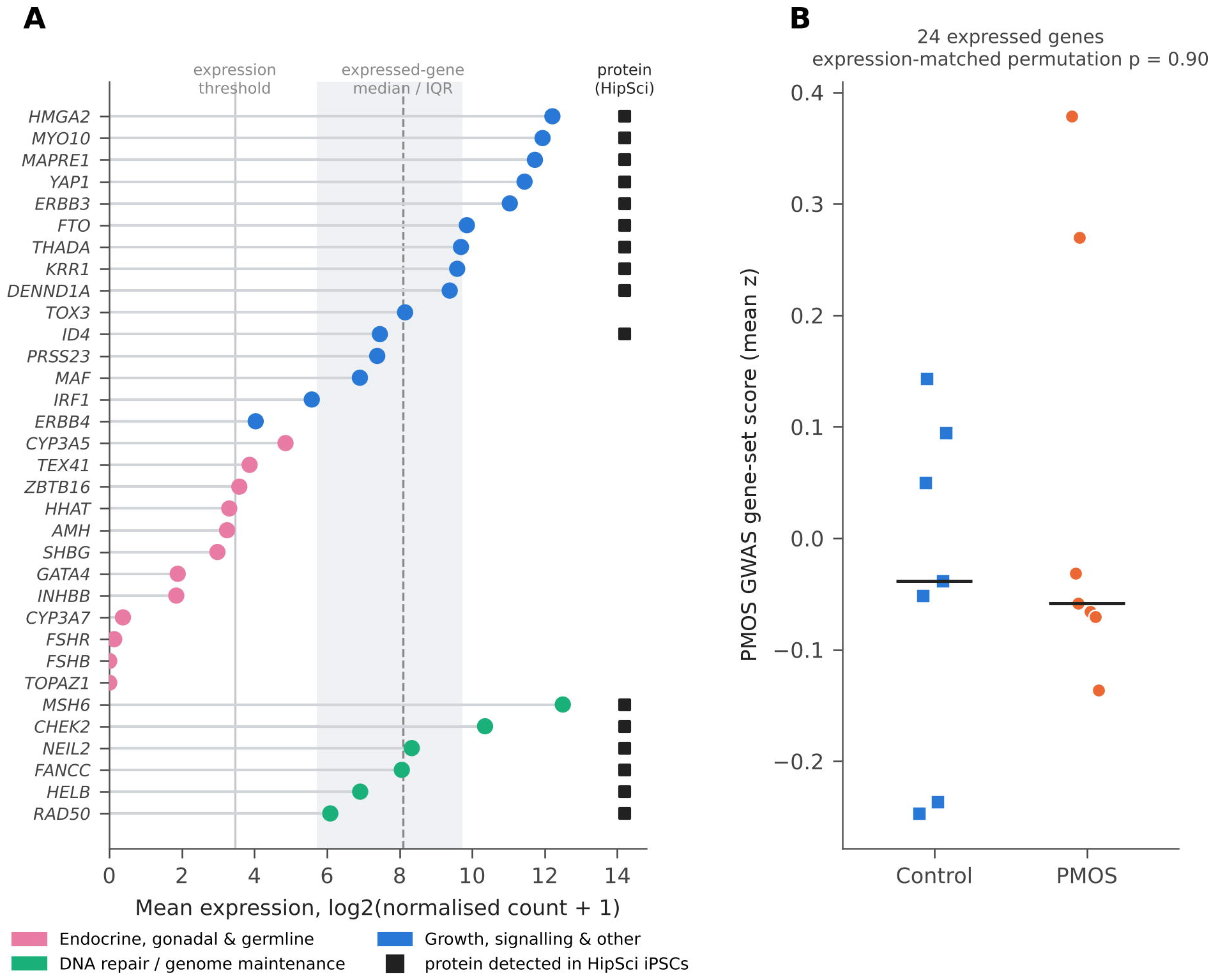
Expression of PMOS GWAS-locus genes in iPSCs. (A) Mean expression of the 33 consensus genes at the 29 PMOS loci (Moolhuijsen et al., 2026), grouped by biological category; black squares, protein detected in HipSci iPSCs (Mirauta et al., 2020). Shaded band, interquartile range of all expressed genes; dashed line, their median; solid grey line, expression threshold. At the “CYP3 complex” locus, for which the GWAS names no gene, CYP3A7 and CYP3A5 are shown. (B) Gene-set score (mean z-score of the 24 expressed genes) by group. p from 10,000 expression-matched random gene sets (primary counts; p = 0.29 with the alternative counts).

A gene-set score of the 24 expressed genes did not differ between groups (Fig. 5B) and was not associated with AMH, FAI, testosterone, BMI or HOMA-IR (expression-matched permutation, all p > 0.29 with either quantification; Additional file 1: Table S6b). At the gene level, THADA and FANCC correlated with FAI (both ρ = 0.62, p = 0.017), and FANCC was slightly higher in PMOS lines (log2FC 0.18, p = 0.049). These nominal findings do not survive multiple testing. In preranked gene-set enrichment analysis, the GWAS consensus genes showed a weak tendency towards higher expression in PMOS lines that did not reach significance consistently (normalised enrichment score +1.4; p = 0.04–0.10 across rankings and calibrations), driven mainly by FANCC, ERBB3, MYO10 and THADA (Additional file 1: Table S6c). With gene-permutation testing, 11–22 Hallmark pathways appeared enriched, but none remained significant after exact sample-label permutation, which preserves gene–gene correlation (Table S6d); there was therefore no pathway-level difference between PMOS and control lines.

Of 60 genes previously linked to PMOS by GWAS or pathobiology, 35 were expressed in iPSCs; only PLGRKT reached nominal p < 0.05 (p = 0.017, q = 1.0; Additional file 1: Table S13), and the set showed no coherent enrichment (Table S6c).

### PMOS-locus genes expressed in iPSCs are largely detected as protein in a reference iPSC proteome

To ask whether the genes expressed in our lines are translated in the pluripotent state, we cross-referenced them with the HipSci iPSC proteome. Overall, 8,726 of the 13,241 expressed protein-coding genes (66%; 53% of all 16,489 expressed genes) are detected in the HipSci iPSC proteome (Mirauta et al., 2020). Detection rose from 5% in the lowest to 96% in the highest expression decile (Additional file 2: Fig. S3A), reflecting the sensitivity of shotgun proteomics rather than absence of protein. Proteins detected in iPSCs were enriched for RNA metabolism, translation, splicing, the mitochondrial matrix, and the cell cycle, whereas expressed genes without detected protein were enriched for G protein-coupled receptors, ion channels, cytokine receptors, and extracellular-matrix proteins (g:SCS-corrected p < 10⁻⁸; Additional file 2: Fig. S3C; Additional file 1: Table S12e). Sixteen of the 24 expressed PMOS consensus genes are detected as proteins in iPSCs: MSH6 (which carries a pQTL in HipSci), CHEK2, RAD50, NEIL2, FANCC, HELB, HMGA2, YAP1, MAPRE1, MYO10, ERBB3, THADA, DENND1A, ID4, KRR1 and FTO (Fig. 5A; Additional file 1: Table S12b). Forty-nine genes combined a nominal PMOS-versus-control difference with detected protein and a reference RNA–protein correlation of at least 0.3 (Additional file 2: Fig. S3B; Additional file 1: Table S12c). None were significant after multiple testing, and the set showed no GO or pathway enrichment, so these genes remain candidates for direct protein measurement.

### PMOS lead variants in the genotyped and imputed data

Seven of the 29 lead variants were on the array; five passed variant QC, whereas rs148982377 was monomorphic, consistent with assay failure, and rs11031005 (FSHB) had a 75% call rate. After imputation, all 29 lead variants were available (5 genotyped, 24 imputed; minimum imputation R² 0.93), with risk-allele frequencies in the 129 QC-passing women close to those of the GWAS (r = 0.984; e.g., rs11075468 0.190 vs 0.184; rs10876864 0.415 vs 0.409; Fig. 6A; Table 2; Additional file 1: Table S3).

**Fig. 6.**
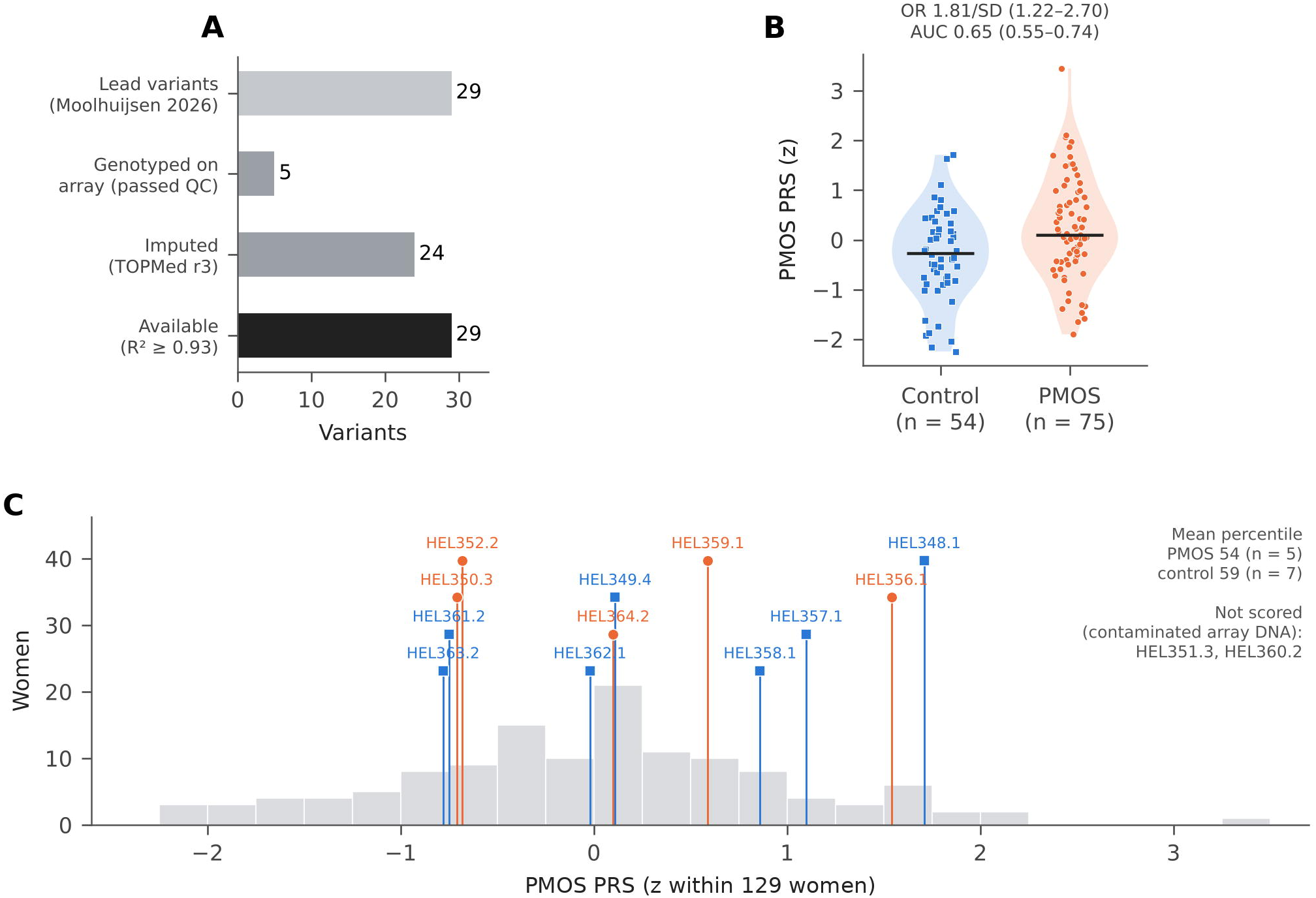
PMOS lead variants and polygenic risk. (A) Availability of the 29 lead variants: 5 genotyped on the array after QC and 24 imputed (TOPMed r3; R² ≥ 0.93), so that all 29 were available. (B) Genome-wide PMOS PRS (PRS-CS, z-score) in women with PMOS (n = 75) and controls (n = 54) among the 129 women passing genotype QC; bars, medians; odds ratio (OR) per SD from logistic regression and AUC with DeLong 95% CI. (C) Distribution of the PRS in the 129 women (grey) with the 12 scored iPSC donors marked (PMOS, orange circles; control, blue squares); HEL351.3 and HEL360.2 were not scored because their array DNA was contaminated.

**Table 2.** PMOS lead variants on the genotyping array. Risk allele and GWAS risk-allele frequency (RAF) from (Moolhuijsen et al., 2026); cohort RAF in the 129 QC-passing women and variant status after QC (Additional file 1: Table S3).

| Variant | Locus (consensus gene) | Risk allele | RAF GWAS / cohort | Status |
|---|---|---|---|---|
| rs10876864 | 12q13 (ERBB3) | G | 0.409 / 0.415 | Genotyped; passed QC |
| rs2070729 | 5q31 (IRF1/RAD50) | C | 0.537 / 0.570 | Genotyped; passed QC |
| rs11075468 | 16q12 (TOX3) | T | 0.184 / 0.190 | Genotyped; passed QC |
| rs8047587 | 16q12 (FTO) | T | 0.452 / 0.388 | Genotyped; passed QC |
| rs1784692 | 11q23 (ZBTB16) | T | 0.840 / 0.845 | Genotyped; passed QC |
| rs11031005 | 11p14 (FSHB) | C | 0.146 / 0.221 | 75% array call rate; imputed (R<sup>2</sup> 0.99) |
| rs148982377 | 7q22 (CYP3 cluster) | T | 0.949 / 0.915 | Monomorphic on the array (assay failure); imputed (R<sup>2</sup> 1.00) |

### Polygenic risk is higher in PMOS across the cohort but does not separate the iPSC donors

Imputation to the TOPMed r3 panel yielded 8,032,962 high-quality SNPs (median imputation R² 0.983), and all 29 lead variants were available (5 genotyped, 24 imputed; minimum R² 0.93). Cohort risk-allele frequencies agreed with those reported for the GWAS (r = 0.984), with concordant effect direction for all 29 variants. The PRS covered 96.8% of the PRS-CS reference panel, compared with 15.2% before imputation. Scores derived with different LD panels and SNP sets were highly concordant (r = 0.94–0.96, compared with 0.71–0.87 before imputation), and restriction to well-imputed SNPs left the score unchanged (r = 0.999).

Across the 129 women passing QC (75 PMOS, 54 controls), the PRS was higher in women with PMOS (mean difference 0.54 SD, 95% CI 0.20–0.87; Wilcoxon p = 0.005; odds ratio 1.81 per SD, 95% CI 1.22–2.70; AUC 0.65, 95% CI 0.55–0.74; Fig. 6B; Table 3; Additional file 1: Table S15). The result was consistent across PRS versions (odds ratio 1.81–1.97 per SD; AUC 0.63–0.67) and sample sets (133 women, AUC 0.64; 128 women without the borderline-contaminated sample, AUC 0.65). The GRS29 pointed in the same direction but more weakly (odds ratio 1.41 per SD, 95% CI 0.98–2.03; AUC 0.61) and correlated only modestly with the genome-wide PRS (r = 0.37), as expected for a score built from the strongest signals alone.

**Table 3.** Genome-wide PMOS polygenic risk of the iPSC donors. PRS (PRS-CS, UK Biobank LD panel) and GRS29 (29 lead variants) as z-score (percentile) within the 129 women passing genotype QC. Cohort comparison (75 PMOS, 54 controls): PRS odds ratio 1.81 per SD (95% CI 1.22– 2.70), AUC 0.65 (0.55–0.74); GRS29 odds ratio 1.41 (0.98–2.03), AUC 0.61 (0.51–0.71). n.s., not scored (array DNA contaminated).

| Line | Group | PRS, z (percentile) | GRS29, z (percentile) |
|---|---|---|---|
| HEL348.1 | Control | 1.71 (96) | 2.00 (97) |
| HEL349.4 | Control | 0.11 (57) | 0.51 (70) |
| HEL357.1 | Control | 1.10 (87) | −0.06 (48) |
| HEL358.1 | Control | 0.86 (83) | 0.18 (57) |
| HEL361.2 | Control | −0.75 (21) | −1.20 (13) |
| HEL362.1 | Control | −0.02 (47) | 0.16 (56) |
| HEL363.2 | Control | −0.78 (19) | 0.14 (55) |
| HEL350.3 | PMOS | −0.71 (22) | 1.82 (94) |
| HEL351.3 | PMOS | n.s. | n.s. |
| HEL352.2 | PMOS | −0.68 (23) | −0.13 (45) |
| HEL356.1 | PMOS | 1.54 (93) | 0.30 (62) |
| HEL359.1 | PMOS | 0.59 (76) | 0.92 (83) |
| HEL360.2 | PMOS | n.s. | n.s. |
| HEL364.2 | PMOS | 0.10 (56) | −0.18 (45) |

Among the 12 scored iPSC donors, PMOS and control donors were spread across the cohort distribution (Fig. 6C; Table 3): the mean PRS percentile was 54 in the five PMOS donors and 59 in the seven control donors. The highest-scoring donor was a control (HEL348.1; 96th percentile for the PRS and 97th for the GRS29), and one PMOS donor (HEL350.3) ranked low on the PRS (22nd percentile) but high on the GRS29 (94th). With five versus seven donors, no group difference could be detected, and the PMOS and control lines should not be regarded as high- and low-risk groups.

## Discussion

In this study, we found no transcriptome-wide evidence of a PMOS-associated expression signature in undifferentiated iPSCs derived from women with a well-defined clinical phenotype. Although the PMOS donors differed from controls in reproductive hormone measures, these clinical differences were not reflected by a coordinated disease-associated transcriptional profile in the pluripotent cells. Instead, expression variation was structured mainly by differences between lines, which in large iPSC collections largely reflect donor genetic background (Rouhani et al., 2014; Kilpinen et al., 2017), although with one clone per donor we cannot fully separate donor from clonal effects; donor age and XIST status made nominal or borderline contributions, whereas diagnosis did not explain more variation than expected under the corresponding null comparisons. The principal implication is therefore specific: under the culture and analysis conditions used here, undifferentiated iPSCs do not provide a robust stand-alone transcriptomic readout of PMOS status. This negative result was robust to an alternative read quantification and to two standard count-based methods, DESeq2 and limma-voom (Law *et al*., 2014; Love *et al*., 2014).

Our findings differ from the extensive differential expression reported in the initial studies of undifferentiated PCOS iPSCs, which compared three affected donors with three controls (Min *et al*., 2018, 2019). Because their full gene lists are not public, only their 20 named top genes could be compared directly: of the 16 expressed in our lines, half changed in the same direction, as expected by chance, and none was significant after correction (Additional file 1: Table S18). Several features may contribute to this difference, and the available evidence does not allow us to identify a single explanation. Small numbers of unrelated donors make estimates sensitive to individual genetic backgrounds and other donor-specific characteristics. Substantial expression differences occur between unrelated iPSC donors even without a disease contrast, and more independent donors make disease-model comparisons more reliable (Germain and Testa, 2017). Limited disease-associated transcriptomic separation has also been reported in patient-derived iPSC models of other complex disorders, and experimental variability and the choice of statistical approach can strongly influence such findings (Valdes *et al*., 2023; Dinasarapu *et al*., 2025). The studies also differed in source tissue, reprogramming method, line characteristics, and analytical strategy (Khatun *et al*., 2023). In particular, the early reports used integrating lentiviral reprogramming and applied nominal significance and fold-change thresholds; the present study used transgene-free CRISPRa-derived lines and evaluated group-level transcriptomic structure using tests that account for the distribution of expression across the panel (Khatun *et al*., 2023, 2026b). These differences mean that the results should not be interpreted as a direct replication under identical conditions. Rather, they show that a large disease-associated signature is not consistently detectable across undifferentiated iPSC panels and underscore the need for adequately powered, independently replicated donor cohorts.

The results also need to be interpreted in the context of cell state. Although some donor-associated molecular characteristics can persist, many regulatory effects are dependent on cell type, differentiation state, or environmental stimulation (Kilpinen *et al*., 2017; Bonder *et al*., 2021). The source eSF phenotype is therefore not expected to be reproduced automatically in undifferentiated iPSCs. The eSF phenotypes described in PCOS (Piltonen *et al*., 2015; Khatun *et al*., 2020, 2021) may depend partly on the hormonal and metabolic environment, may involve epigenetic features that are modified during reprogramming, or may require a differentiated, hormone-responsive cell state. Our data cannot distinguish these possibilities. They do, however, support testing the same donor lines after directed differentiation into relevant cell types, including endometrial stromal, granulosa-like, or other ovarian-supporting lineages, followed by appropriate hormonal or metabolic challenges (Khatun *et al*., 2023, 2026b). The recent demonstration of androgen-associated dysfunction in iPSC-derived endothelial cells from women with phenotype A PCOS further illustrates how a disease-related phenotype may become apparent in a differentiated cellular context (Wu *et al*., 2026).

The expression of genes at PMOS-associated loci provides a further indication that the pluripotent state captures only part of the biology implicated by genetic association. Several genes involved in reproductive hormone signalling or endocrine regulation, including FSHR, FSHB, AMH, SHBG, INHBB and GATA4, were not detectably expressed in the undifferentiated cells under the conditions examined. Regulatory effects involving genes that are inactive or expressed at very low levels in this state are unlikely to be adequately assessed using these cells without differentiation or an appropriate stimulation protocol. In contrast, a number of genes involved in DNA damage responses, genome maintenance and growth regulation— including MSH6, CHEK2, RAD50, NEIL2, FANCC, HELB, HMGA2, YAP1, ERBB3, THADA and DENND1A—were expressed (Moolhuijsen *et al*., 2026). This observation identifies potentially accessible molecular pathways for follow-up, but expression alone does not establish that these genes mediate PMOS risk in iPSCs or in the ovary. Similarly, associations between PMOS loci, DNA-damage-response pathways and reproductive ageing should not be taken as direct evidence that the present pluripotent cells model ovarian reserve or oocyte longevity. Such hypotheses require testing in appropriately specified germ-cell or ovarian cell models.

Human primordial germ-cell-like cells (hPGCLCs) offer one route to early germline specification, but hPGCLC identity and induction do not equate to the formation of functional oocytes or the recapitulation of ovarian reserve. Protocols and efficiencies remain variable, and interpretation requires careful validation of cell identity and developmental state (Vaz Santos *et al*., 2025). Applying a validated hPGCLC differentiation strategy to the present donor panel could test whether selected PMOS-associated genetic effects emerge during early germline specification, but it would not, by itself, establish a mechanism for ovarian ageing or fertility outcomes.

XIST expression identified an additional source of heterogeneity relevant to the interpretation of female iPSC models. Two of the 14 lines had clearly lost XIST expression and a third (HEL363.2) was borderline, and these lines also showed evidence consistent with altered expression of the X-linked gene KDM5C. The finding is potentially indicative of XCI erosion (Mekhoubad *et al*., 2012; Vallot *et al*., 2015), which is heterogeneous between lines and can persist after differentiation (Thapa *et al*., 2026). In our panel, the distribution of low-XIST lines was uneven between diagnostic groups, making XIST status a potential confounder in case–control analyses. Nevertheless, XIST expression alone is an indirect marker: it does not establish the extent of chromosome-wide reactivation, nor does it define the functional consequences in each line. Its quantification also depends on the counting method: XIST overlaps the antisense TSIX transcript, so unstranded counting of a stranded library splits reads between the two genes and underestimates XIST, which in our data determined whether HEL363.2 fell below the erosion cut-off. Studies that classify lines by XIST should therefore report the counting mode and the threshold used. Follow-up assessment could include additional X-linked gene-expression markers, allele-specific expression and, where appropriate, DNA-methylation or chromatin-based assays.

Genotyping the donors within a larger cohort was intended to answer a question that 14 lines alone cannot: did selecting donors by clinical diagnosis also select them for inherited PMOS liability? Two observations address this. First, across 129 women from the same clinical population, the genome-wide PRS was higher in women with PMOS (odds ratio 1.81 per SD; AUC 0.65), a modest discrimination consistent with that reported for PCOS polygenic scores in other cohorts (Joo *et al*., 2020; Moolhuijsen *et al*., 2026); suggesting that the score captures part of PMOS liability in this Finnish setting. Second, despite this, the PMOS and control iPSC donors overlapped completely in the cohort distribution, and the highest-scoring donor was a control. The panel therefore contrasted diagnoses, not genetic liabilities. This offers a possible genetic explanation, alongside cell state, for the absence of a PMOS transcriptomic signature: if the cellular effects of PMOS risk are polygenic and small per allele, a panel without a contrast in polygenic load is not expected to reveal them. An important practical implication is that future PMOS iPSC panels should be assembled genotype-first, by selecting donors from genotyped cohorts at the extremes of polygenic risk and, ideally, concordant with diagnosis (high-PRS cases versus low-PRS controls), as is now done for other polygenic disorders (Page *et al*., 2022; Sawada *et al*., 2024). The genotyped cohort described here can serve as the sampling frame for such recruitment.

The polygenic-risk analyses also require cautious interpretation. The cohort comparison was descriptive and was not adjusted for all potential sources of confounding, including genetic principal components. The Oulu cohort is independent of the PMOSGWAS: none of its participants contributed to the discovery samples (Moolhuijsen *et al*., 2026). The case-control association of the score is therefore an out-of-sample estimate and is not inflated by sample overlap, a common source of bias in PRS analysis (Choi *et al*., 2020). However, PRS accuracy declines with genetic distance from the discovery population (Martin *et al*., 2018), and the Finnish population, although of European ancestry, has a distinct genetic history. The present sample is too small to establish whether polygenic score is a better predictor of iPSC expression than diagnosis, and the score should not be interpreted as a clinical diagnostic tool in this study.

The reference proteome adds a protein-level dimension to these genetic observations, but its scope must be stated precisely. In HipSci, mRNA and protein abundance were only partly correlated across iPSC lines, and some disease-linked pQTLs were detectable only at the protein level (Mirauta *et al*., 2020), as expected from the post-transcriptional control of protein levels (Liu *et al*., 2016). Against this reference, 16 of the 24 PMOS-locus genes expressed in our lines, including the DNA-repair and growth genes, are translated into detectable protein in undifferentiated iPSCs, and MSH6 carries a cis-pQTL. These genes are therefore candidates whose genetic regulation could be studied at the protein level in the pluripotent state, whereas the endocrine locus genes, absent at the RNA level, are also absent from the iPSC proteome and require differentiated models. The proteome does not uncover a hidden disease signature, and direct proteomic profiling of genotype-selected lines will be required to test protein-level effects.

Taken together, the results have practical implications for the design of iPSC models of complex endocrine and reproductive disorders. First, the independent donor—not the number of wells or technical replicates—must remain the biological unit for disease-group inference. Donor numbers should be justified prospectively, with sufficient unrelated individuals to reduce the influence of background variation (Germain and Testa, 2017). Second, the source tissue, reprogramming method, passage history, culture conditions, and genomic characteristics of each line should be reported and considered in interpretation. Third, female lines should be assessed for XCI status, with XIST expression treated as a screening indicator rather than a definitive standalone assay. Fourth, the choice of cellular state should follow the biological question: undifferentiated iPSCs are useful for studying pluripotency-associated and broadly expressed pathways, whereas endocrine, ovarian and endometrial mechanisms may require differentiation and defined hormonal or metabolic exposures. These principles are consistent with current efforts to improve the diversity, transparency and interpretability of human pluripotent stem-cell model systems (Khatun *et al*., 2026b).

## Strengths and limitations

This study has several strengths. It examines 14 clonal iPSC lines from seven PMOS and seven control donors, more independent donors than previous transcriptomic comparisons of PMOS iPSCs, which included three or four donors per group. The lines were derived from eSF, generated using CRISPRa-based transgene-free reprogramming and characterised as described previously (Khatun *et al*., 2026a). The patient group was clinically well defined, with all seven donors meeting all three Rotterdam criteria. The study also combines transcriptomic data with detailed donor phenotyping and genome-wide genotype information and explicitly examines XIST expression as a potential source of variation. Gene-set and pathway analyses were calibrated by exact sample-label permutation, which avoids the false-positive enrichment that gene-permutation tests produce in small panels. The use of panel-level exact and permutation-based analyses provides an additional way to assess whether observed group structure exceeds that expected from alternative assignments of the donors. Finally, the transcriptome results were robust to an alternative read quantification and to count-based differential-expression methods (DESeq2 and limma-voom).

Several limitations should be considered. First, despite being larger than previous PMOS iPSC comparisons, 14 lines provide limited power to detect modest expression effects expected for a polygenic condition. The absence of transcriptome-wide significant differences should therefore not be interpreted as proof that all disease-associated effects are absent; it indicates that large, consistent effects were not detected under the conditions studied. Second, one clone per donor was analysed, so donor and clone effects cannot be formally separated; attributing the line-level variation mainly to donor background rests on evidence from larger iPSC collections (Rouhani et al., 2014; Kilpinen et al., 2017). Third, only undifferentiated iPSCs were profiled, and the results cannot be extrapolated directly to differentiated endometrial, ovarian, endothelial, or germline cells. Fourth, controls were frequency-matched to cases by BMI category rather than individually matched, and median BMI remained higher in PMOS donors (31.1 vs 23.4 kg/m²); although BMI-category and continuous-BMI adjustment did not change the result, residual confounding by BMI or correlated metabolic traits cannot be excluded with 14 donors. Fifth, the presence of a mosaic idic(20) in HEL350.3 and contaminated array DNA for HEL351.3, as described in the line-characterisation study, may affect some line-or genotype-level analyses; the RNA-seq of HEL350.3 shows the expected chromosome 20 dosage effect, so its chromosome 20 genes should be interpreted with caution, although the reported sensitivity analyses did not materially alter the main transcriptomic findings. Sixth, the protein comparison relied on a reference proteome from other iPSC donors rather than proteomic measurements in the 14 lines studied here. Accordingly, protein-level observations provide context but cannot establish protein abundance in these specific lines. Seventh, nominal gene-level results, including the top-ranked genes and the XIST status of HEL363.2, depended on read quantification (stranded versus unstranded counting and the handling of multi-mapping reads), although the transcriptome-wide conclusions did not. Finally, only 12 of the 14 iPSC donors could be scored for polygenic risk. These limitations reinforce the exploratory status of the genetic association analyses.

## Conclusions

In conclusion, undifferentiated iPSCs derived from women with clinically defined PMOS showed no robust diagnosis-associated transcriptomic differences in this 14-line study. This finding does not rule out inherited PMOS mechanisms; rather, it indicates that such mechanisms may not be expressed uniformly in the pluripotent state, and disease-associated differences may emerge only in differentiated, hormone-responsive cells. Donor-associated line-to-line variation and XIST status are important considerations when interpreting female iPSC transcriptomes, while the expression of selected PMOS-locus genes suggests that the pluripotent state may be informative for some pathways but not others. Diagnosis-based donor selection did not create a contrast in polygenic risk, so donors should be selected by both phenotype and polygenic risk. Future work should combine adequately sized, genotyped donor panels with validated differentiation into disease-relevant cell types and carefully controlled hormonal or metabolic exposures (Khatun *et al*., 2026c). Such an approach will provide a more appropriate framework for testing how PMOS-associated genetic variation contributes to reproductive and metabolic phenotypes.

## List of abbreviations

AMH: anti-Müllerian hormone
AUC: area under the receiver operating characteristic curve
BMI: body mass index
CRISPRa: CRISPR activation
eQTL: expression quantitative trait locus
eSF: endometrial stromal fibroblasts
FAI: free androgen index
FDR: false discovery rate
FG: Ferriman–Gallwey
GO: gene ontology
GRS29: 29-variant genetic risk score
GSA: Global Screening Array
GWAS: genome-wide association study
HOMA-IR: homeostasis model assessment of insulin resistance
iPSC: induced pluripotent stem cell
LD: linkage disequilibrium
MAF: minor allele frequency
OR: odds ratio
PCA: principal component analysis
PCOS: polycystic ovary syndrome
PMOS: polyendocrine metabolic ovarian syndrome
pQTL: protein quantitative trait locus
PRS: polygenic risk score
QC: quality control
RAF: risk-allele frequency
SD: standard deviation
STR: short tandem repeat
TOPMed: Trans-Omics for Precision Medicine
XIST: X-inactive specific transcript.

## Declarations

### Ethics approval and consent to participate

The study was approved by the Regional Medical Research Ethics Committee of the Wellbeing Services County of North Ostrobothnia (EETTMK 58/2026). All participants gave written informed consent in accordance with the Declaration of Helsinki.

### Consent for publication

Not applicable.

### Availability of data and materials

RNA-seq data will be deposited in the Gene Expression Omnibus (GEO) or, where required by the consent conditions, a controlled-access repository, and the accession number will be provided when available. Both the primary and the reverse-stranded count matrices will be provided. Individual-level genotypes cannot be shared openly under the ethics approval; access is available on request. The iPSC lines are registered in hPSCreg (UHi023-A to UHi036-A). Analysis code (release 1.0.1) is summarized in Additional file 3 and is available at GitHub (https://github.com/BrayanRodriguez2/PMOS-iPSC-study-analysis-code) and Zenodo (https://doi.org/10.5281/zenodo.24025021, concept DOI), with code under the MIT licence and aggregated result tables under CC BY 4.0. Donor-level data (genotypes, clinical data, per-donor scores and counts) are not included in the release and are available on reasonable request through controlled access. The HipSci reference data are publicly available with reference (Mirauta et al., 2020).

## Authors’ contributions

MK conceived and designed the study, secured funding, established and characterised the iPSC lines, performed the experiments, carried out the transcriptomic analyses (differential expression, variance and sensitivity analyses, and the HipSci reference-proteome comparison), prepared the figures and tables, and wrote the manuscript. BSRM performed the genetic analyses (genotype quality control, imputation, and polygenic risk scoring) and the RNA-seq quantification and sensitivity analyses, including gene-set enrichment and comparison with published iPSC signatures, and prepared the code release. AA performed cDNA synthesis and contributed to RNA-seq analysis. VM supervised the data analysis. KL and TT contributed expertise in stem cell biology. RKA and TTP recruited and phenotyped the donors. JST and AS supervised the study and provided resources. All authors critically revised the manuscript and approved the final version.

## Use of generative AI

Claude (Anthropic; Claude Opus, accessed June-November 2026) was used to write and run analysis code for the transcriptomic and protein-reference analyses, for language editing, and for editing labels and layout of the final figures. All analyses used the authors’ own data. All code was independently revalidated by BSRM, and every reported value was checked against the original result files by the authors. The authors take full responsibility for the content of this article.

### Additional files

**Additional file 1 (.xlsx):** Supplementary Tables S1–S18, one sheet per table, with a README sheet (contents listed below).

**Additional file 2 (.pdf):** Supplementary Figs. S1–S3 with legends.

**Additional file 3 (.docx):** Computational methods and code availability: a summary of the computational methods, software and random seeds, and a map from each figure and table to its script. The code (release 1.0.1) is available at GitHub and archived at Zenodo (https://doi.org/10.5281/zenodo.24025021).

**Additional file 2: Supplementary Figs. S1–S3.** Fig. S1. (A) PCA of all 16,489 expressed genes; colour, diagnosis; shape, BMI category. (B) Spearman correlation of PC1–PC5 (1,000 most variable genes) with donor traits and XIST; asterisk, p < 0.05 (nominal; no correlation remains significant after correction for the 40 tests). (C) Distribution of gene-wise p-values for PMOS versus control, without (bars) and with (line) XIST as a covariate; dashed line, uniform expectation. Fig. S2. Z-scored expression of the top 40 genes by nominal p-value (PMOS versus control) in all 14 lines; none reaches FDR < 0.1; illustrative only, as the ranking depends on quantification (25 of the 40 genes remain in the top 40 with the alternative quantification). Fig. S3. Protein-level cross-reference with the HipSci iPSC proteome (Mirauta et al., 2020). (A) Percentage of expressed protein-coding genes (bars) and of all expressed genes (black ticks) with protein detected in HipSci, by expression decile in our lines. (B) Z-scored mRNA expression of the 49 genes with nominal PMOS-versus-control p < 0.05, detected protein, and reference RNA–protein correlation r ≥ 0.3 (r after each gene). (C) Selected GO and pathway terms enriched among expressed genes with (blue) or without (grey) detected protein, relative to all expressed protein-coding genes (g:Profiler; Kolberg et al., 2023; g:SCS-corrected p).

| Table | Content |
|---|---|
| S1 | Donor clinical characteristics, hPSCreg identifiers and XIST status (n = 14) |
| S2 | Genotype QC summary: numbers of samples passing and failing each criterion (call rate, heterozygosity, sex, relatedness, contamination) and final counts (n = 140); per-sample records are held by the authors |
| S3 | The 29 PMOS lead variants (Moolhuijsen et al., 2026) with array availability, imputation status and R<sup>2</sup>, strand, and RAF in the 129 QC-passing women |
| S4 | Gene-wise PMOS vs control results, with and without XIST adjustment (16,489 genes) |
| S5a–c | Trait association summary, transcriptome-wide permutation tests, gene–trait associations at p < 0.01 |
| S6a–d | Expression of 33 PMOS-locus consensus genes, gene-set score tests, preranked GSEA of GWAS and literature genes, and Hallmark GSEA with exact label-permutation calibration |
| Table | Content |
| S7 | Sensitivity analyses excluding HEL350.3 and HEL351.3, HEL350.3 only, or the two XIST-eroded lines with and without the borderline HEL363.2 |
| S8 | Sample-to-sample correlation matrix and within/between-group tests |
| S9 | Variance partition per gene (PMOS, BMI, age, XIST) with permutation null |
| S10 | BMI-adjusted and PMOS × BMI interaction models (16,489 genes) |
| S11 | Donor characteristics within BMI strata (BMI <25 and >30 kg/m<sup>2</sup>) |
| S12 | Protein-level cross-reference with the HipSci iPSC proteome (original HipSci data): detection by expression, PMOS-locus genes, 49 candidates, GO enrichment, all genes |
| S13 | 60 literature-nominated PMOS genes in the gene-wise results |
| S14 | Top 40 nominal genes with functional annotation (illustrative; not robust to quantification) |
| Table | Content |
| S15 | Imputation and PRS summary: SNP coverage, agreement between PRS versions, and case–control results for each PRS version and sample set |
| S16 | Detectable fold change by number of donors per group (80% power; per-gene α = 0.05 and 0.001), based on the between-line variance of this panel |
| S17 | Sensitivity analyses: alternative (reverse-stranded) quantification, library QC, agreement of count matrices, DESeq2 and limma-voom models, XIST-dependent results, chromosome 20 dosage in HEL350.3 |
| S18 | The 20 named top genes of Min et al. (2018) in the gene-wise results (direction and significance) |

